# A comprehensive *Drosophila* toolkit reveals context-dependent functional hierarchies across Ras^G12X^ variants *in vivo*

**DOI:** 10.64898/2026.11.19.767789

**Authors:** Miriam Rodríguez-Vázquez, Maria-Armineh Tossounian, Nsikan Nsek, Matthew B Ames, Isabel Liseth, Andrea Foley, Susumu Hirabayashi, Helena M Cochemé

**Affiliations:** MRC Laboratory of Medical Sciences (LMS), Du Cane Road, London W12 0HS, UK; Institute of Clinical Sciences, Imperial College London, Hammersmith Hospital Campus, Du Cane Road, London W12 0HS, UK; Department of Cancer Metabolomics, Sasaki Institute, Sasaki Foundation, 2-2 Kandasurugadai, Chiyoda-ku, Tokyo 101-0062, Japan

**Keywords:** KRAS, Ras signalling, G12X mutations, G12X variants, oncogenes, *Drosophila*, development, cancer models, *in vivo* transgenic toolkit

## Abstract

KRAS is a GTPase regulating key signalling processes involved in cell proliferation, differentiation, growth and survival. KRAS is the most prevalent oncogene in human cancers, often acquiring a point mutation at Glycine 12 (G12). Although several possible amino acid substitutions occur at this hotspot in humans, most animal cancer models often examine only a single or limited subset of G12X variants, restricting our understanding of their distinct biological effects. Here, we describe a novel transgenic toolkit in *Drosophila melanogaster* comparing the most frequently observed oncogenic G12X variants (G12A, G12C, G12D, G12R, G12S, G12V), and evaluate these across multiple developmental and tumour contexts *in vivo*. Interestingly, we demonstrate how the phenotypic strength of each G12X variant differs significantly depending on the biological context, highlighting the importance of considering these point mutations systematically. Overall, our comprehensive Ras^G12X^ toolkit provides a powerful and versatile platform to explore the dysregulation of KRAS *in vivo*, serving as a valuable resource enabling unprecedented opportunities to investigate variant- and context-specific mechanisms of oncogenic signalling with direct relevance to cancer biology and therapy.

## Introduction

The Ras family of GTPase enzymes regulates key cellular processes, including proliferation, differentiation and survival^1,2^. Ras acts as a molecular switch, cycling between active GTP-bound and inactive GDP-bound states (Fig. 1A). Ras controls several major downstream signalling pathways, including RAF/ERK/MAPK and PI3K/AKT. Dysregulation of Ras signalling leads to a range of developmental disorders, known as RASopathies^3^. Furthermore, Ras is amongst the most frequently mutated genes in human cancers^4^. Indeed, activating mutations in KRAS occur in ∼41% of colon adenocarcinomas and ∼86% of pancreatic adenocarcinomas^5^.

**Figure 1.**
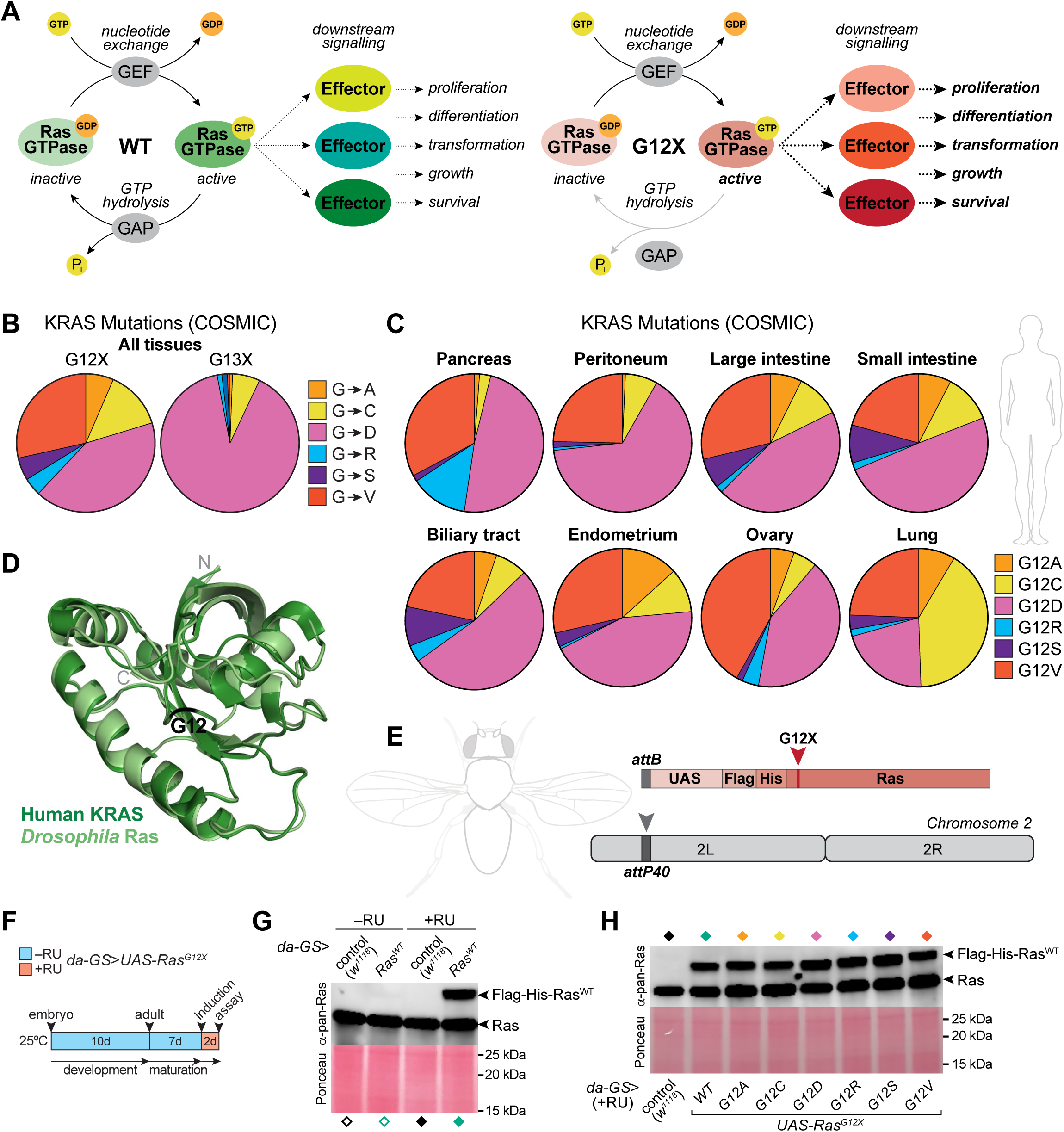
Generation and validation of the Ras^G12X^ *Drosophila* toolkit. (**A**) The Ras GTPase cycles between a GDP-bound inactive and a GTP-bound active form. GEFs (<u>g</u>uanine nucleotide <u>e</u>xchange factor) promote GDP-to-GTP exchange switching Ras on, while GAPs (<u>G</u>TPase-<u>a</u>ctivating <u>p</u>rotein) accelerate GTP hydrolysis turning Ras off. Mutation at G12X prevents interaction with GAP, therefore locking Ras in a constitutively active state. (**B**) Relative frequency of KRAS^G12X^ and KRAS^G13X^ missense mutations in human cancers from the COSMIC database for all tissues (see Table S2). (**C**) Distribution of KRAS^G12X^ variants in human cancers across tissue types (see Tables S3 and S4). (**D**) Superposition of the human KRAS structure (dark green; PDB: 4OBE, residues 1-169) with the AlphaFold 3 model of *Drosophila* Ras (pale green; residues 1-169). RMSD (root mean square deviation) = 0.378 Å for 138 atoms. G12 is indicated in black. See Fig. S1B for the individual structures. (**E**) Map of the *UAS-Ras^G12X^*transgenes (including N-terminal Flag and His tags), inserted at the attP40 site of chromosome 2. (**F**) Experimental scheme showing the generation of adults expressing Ras^G12X^ ubiquitously (left). Developmental effects were bypassed using the drug-inducible GeneSwitch system. Female flies were analysed after 48 h of induction from day 7 of adulthood. (**G**,**H**) Western blotting with pan-Ras antibody detecting endogenous Ras (lower band) and the tagged transgenic protein (upper band). Confirmation that the *UAS-Ras^G12WT^* transgene is expressed only in the presence of the inducer drug RU (**G**), and that the *UAS-Ras^G12X^*variants are expressed at similar levels (**H**). Ponceau staining is shown as a loading control.

According to the COSMIC (<u>C</u>atalogue <u>o</u>f <u>S</u>omatic <u>M</u>utations in <u>C</u>ancer) database of human tumour tissues, simple coding mutations within the KRAS gene are identified in 49,216 out of 253,224 samples tested^6^. These KRAS mutations consist predominantly (>99%) of missense substitutions (Table S1). Strikingly, ∼84% of these point mutations occur at Glycine 12 (G12). Several possible amino acid substitutions are detected at this hotspot, including G12A, G12C, G12D, G12R, G12S and G12V. Interestingly, these KRAS^G12X^ variants are present at different frequencies (Fig. 1B, Table S2). Across all tissues, the G12D variant is the most common (∼42%), followed by G12V (∼28%) and G12C (∼14%). This pattern contrasts with the G13 site, representing ∼14% of KRAS point mutations, where ∼90% of substitutions are G13D (Fig. 1B, Table S2). Importantly, the distribution of the KRAS^G12X^ variants differs considerably between tumour tissue types. While G12D tends to be the most common variant (e.g. ∼65% in peritoneal, ∼49% in pancreatic), the G12C mutation is strongly enriched in lung cancer (∼41%) (Fig. 1C, Tables S3,S4). This prevalence likely arises from variant- and tissue-specific genetic interactions, highlighting considerable context-dependent complexity underlying their patterns of observed frequency^5,7^.

Given its central role in cellular physiology and its dysregulation in diseases such as cancer, KRAS has been studied extensively. However, despite reported differences in the oncogenic properties of G12X variants *in vitro*^8–11^, animal studies have generally examined only a single variant or limited subset of variants (e.g. ^9,12–15^). Consequently, research to-date provides only a partial view of how variant identity and tissue context shape Ras signalling and biological outcomes. The functional and physiological differences between the G12X variants therefore remain incompletely characterised *in vivo*, which limits our fundamental biological understanding of their context-specific effects.

*Drosophila melanogaster* is an important experimental model system to study development, metabolism, cancer and ageing^16–26^. Ras proteins in *Drosophila* are represented by Ras1 (Ras85D; CG9375) and Ras2 (Ras64B; CG1167). Ras1 corresponds to mammalian KRAS/HRAS/NRAS, while Ras2 is equivalent to mammalian RRAS. In this study, we refer to Ras1 as Ras, in accordance with the convention prevalent in the field. *Drosophila* Ras and human KRAS exhibit strong evolutionarily conservation, as highlighted by sequence alignment (∼79.8% identity for the full protein, ∼93.3% identity across residues 1-120; Fig. S1A) and structural modelling (Figs. 1D and S1B). Notably, the first 79 amino acids are 100% identical between flies and humans, which encompass important functional regions: the ‘P-loop’ motif (responsible for phosphate binding), the ‘Switch I’ region (forming the primary binding interface for downstream effector proteins such as RAF and PI3K, and regulators such as GAPs and GEFs), and the ‘Switch II’ region (containing a conserved catalytic glutamine, Q61, required for GTP hydrolysis) (Figs. 1A and S1A,B).

Oncogenic activation of *Drosophila* Ras causes developmental defects and tissue hyperplasia, mimicking aspects of cancer^27–31^. Historically, the Ras^G12V^ variant is widely used in the *Drosophila* field as the stereotypical activating mutant of Ras signalling, and has contributed to many important discoveries in the areas of developmental growth and cell biology^27–31^. In contrast, the other G12X mutations remain largely uncharacterised, and the available reagents are considerably more limited (e.g. ^32^). To address this knowledge gap, we leverage the powerful and tractable *Drosophila* system to generate a comprehensive toolkit of Ras^G12X^ variants, allowing their detailed and comparative profiling at an organismal level and in tissue-specific contexts. This resource has enabled us to study the impact of different Ras^G12X^ variants in a range of developmental and proliferative settings, revealing striking context-dependent differences in their relative functional strength and phenotypic effects *in vivo*.

## Results

### Generation & validation of a novel Ras^G12X^ transgenic toolkit

We generated a comprehensive *Drosophila* toolkit of UAS transgenic lines for the major KRAS^G12X^ variants described in human tumours: G12A, G12C, G12D, G12R, G12S and G12V. All lines were created by targeted insertion at the same *attP40* docking site via *ɸC31* integrase, thereby ensuring consistent expression between transgenes, allowing for reliable comparative studies (Fig. 1E). The *UAS-Ras^WT^* and *UAS-Ras^G12X^* lines contain N-terminal Flag and His tags, facilitating downstream molecular analysis (e.g. purification, immunoblotting). Presence of the correct substitution was confirmed by sequencing analysis (Fig. S1C, Table S5).

Since ubiquitous constitutively activated Ras^G12X^ variants are developmentally lethal, we used the drug-inducible GeneSwitch (GS) system^33^ to achieve adult-specific transgene expression. Flies were allowed to develop and mature in the absence of the inducer drug RU (RU486/mifepristone), then switched onto +RU food at 7 days of adulthood for 48 h of induction (Fig. 1F). We assessed protein abundance by immunoblotting using a pan-Ras antibody (upper band=tagged Ras^G12X^ variant, lower band=untagged endogenous Ras). We confirmed that the *UAS-Ras^WT^* line was only expressed in the presence of RU drug, and did not affect levels of endogenous Ras (Figs. 1G and S1D). Next, we assessed the full G12X collection. Protein levels between the Ras^G12X^ variants were consistent (Figs. 1H and S1E), supporting that the mutations do not appear to affect protein stability or turnover. Overall, we have demonstrated that our novel comprehensive toolkit is appropriate for functional analysis of the Ras^G12X^ variants *in vivo*. As a benchmark, we have also included experiments with the existing widely used original G12V line, *UAS-Ras^G12V^*(P) (=*UAS-Ras1^V^*^12^), generated by random P-element insertion^28^.

With the above validated toolkit, we can exploit the powerful genetics of the *Drosophila* system to functionally compare these Ras^G12X^ variants in a range of physiological contexts. Here, we effectively leverage the fly as an *in vivo* biosensor to rapidly gain critical insights into how the Ras^G12X^ mutations determine cellular differentiation and tissue growth, both during development and in tumour models.

### Effect of Ras^G12X^ on development (1): Developmental timing, pupariation rates, viability and survival

As an initial characterisation of our toolkit, we expressed the *UAS-Ras^G12X^*transgenes in different contexts related to development. First, to test how the G12X variants impact development and viability at a whole organism scale, we used the ubiquitous driver *da-GAL4* (*daughterless*) to express the *UAS-Ras^G12X^* lines constitutively (Fig. 2A). Expression of wild-type (WT) Ras (*da-GAL4*>*UAS*-*Ras^WT^*) did not affect pupariation, body size or developmental timing compared to control (*da-GAL4>w*^1118^), indicating that overexpression of WT Ras at this level does not perturb normal physiological development (Fig. 2B-E).

**Figure 2.**
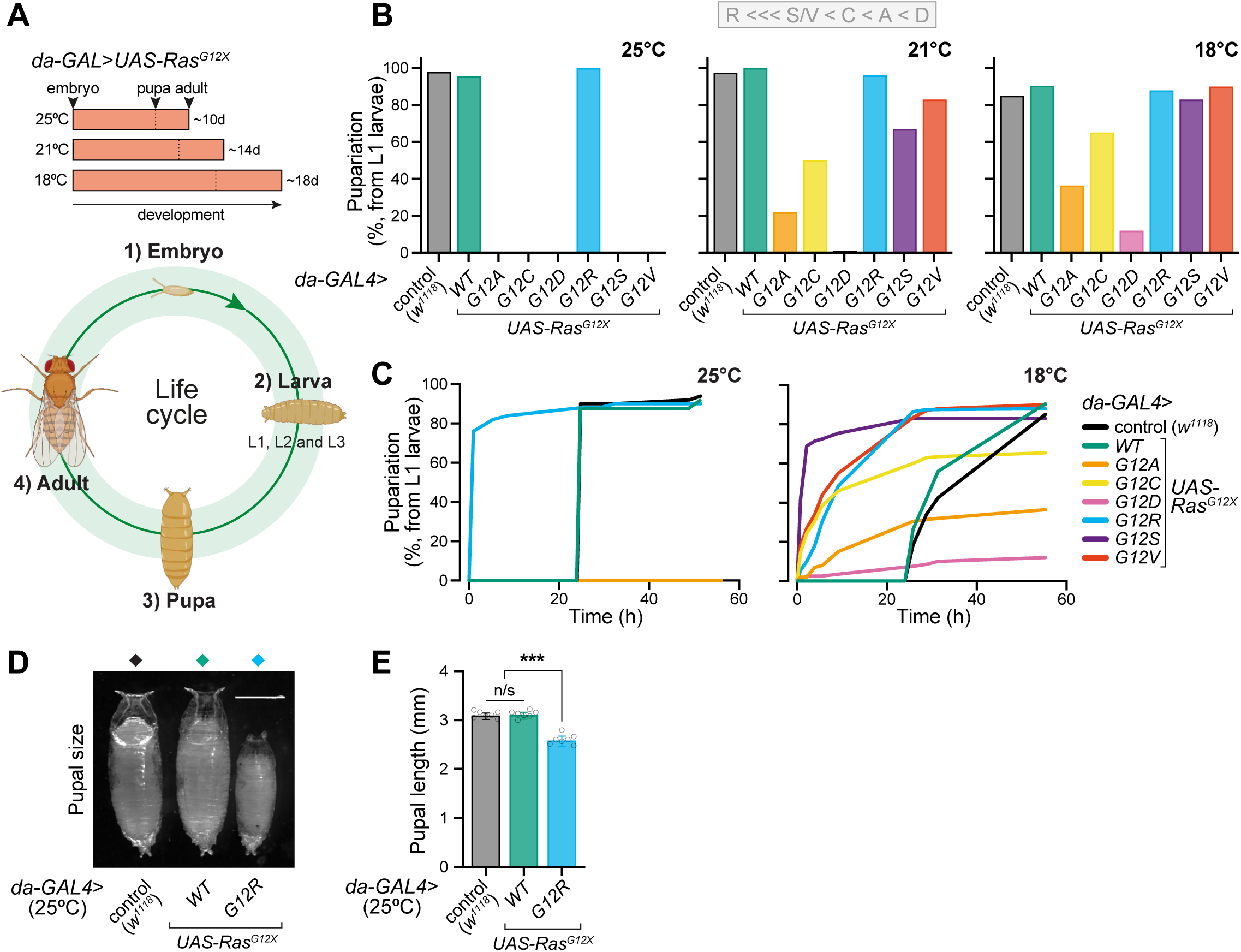
Whole body expression of Ras^G12X^ variants differentially affects development and pupariation. (**A**) Experimental scheme with approximate developmental timings of flies at 25°C, 21°C and 18°C. The *Drosophila* graphics for the lifecycle were obtained from BioRender. (**B**,**C**) Pupariation rate (% of L1 larvae picked) for *da-GAL4>UAS-Ras^G12X^* flies at 25°C, 21°C and 18°C (**B**). Developmental timecourse of pupariation at 25°C and 18°C (**C**). Experiments were set up with a total of n=50 (25°C) or n=200 (21°C and 18°C) L1 larvae. (**D**,**E**) Representative imaging of pupal size at 25°C (scale bar: 1 mm), for the controls (*da*>*w*^1118^ and *da*>*Ras^G12WT^*) and the *da>Ras^G12R^* variant (**D**). Quantification of pupal length (**E**). Data are means ±SD of n=7 replicate, analysed by one-way ANOVA. n/s, p>0.05; ***, p<0.001.

Incubating the developing flies at different temperatures (18°C, 21°C or 25°C) modulates the level of transgene expression, which increases as temperature rises. Therefore, temperature can act as a rheostat for the Ras signalling pathway, allowing us to achieve finer resolution in our phenotypic stratification. For instance, from picked L1 larvae, out of all G12X mutants, only G12R pupariated at 25°C, whereas at 21°C all the G12X variants except G12D formed pupae to varying extents, with G12A and G12C appearing more severe than G12S and G12V (Fig. 2B,C). The original P-element *Ras^G12V^*(P) line behaved more strongly than the toolkit attP40 *Ras^G12V^* (Fig. S2A). Note that none of the *da-GAL4*>*UAS-Ras^G12X^*pupae gave rise to viable adults (Fig. S2B), and these were considerably smaller than WT pupae (Fig. 2D,E), presumably due to effects on hormonal signalling, as previously reported for the G12V mutant^34^. In addition to effects on body size, we also recorded developmental timing, and observed that the G12R mutant at 25°C pupariated more rapidly than the overlapping *w*^1118^ and Ras^WT^ controls (accelerated by ∼24h, Fig. 2C). Therefore, from these whole body *in vivo* readouts, we can already uncover clear differences between the Ras^G12X^ variants. Next, we applied an analogous approach focussing on two specific tissues that are highly amenable and frequently used as outputs of large-scale phenotypic screens in *Drosophila*: the eye and the wing.

### Effect of Ras^G12X^ on development (2): Eye developmental phenotyping

The *Drosophila* compound eye is composed of ∼750 ommatidial units, each assembled from multiple photoreceptor, cone and accessory cells. The adult eye develops from the larval eye imaginal disc (undifferentiated pluripotent cells), with the ommatidia arranged into a highly ordered pattern^35^. Disruption of the Ras signalling cascade during eye development results in a so-called ‘rough eye’ phenotype from disorganisation of the ommatidial array, as well as structural deformations. We drove expression of the *UAS-Ras^G12X^* variants in the developing eye using the *GMR-GAL4* driver, and scored effects on the severity of the resulting phenotype at a range of temperatures.

As expected, expression of the original P-element *UAS-Ras^G12V^*(P) line was predominantly pupal lethal at 25°C^28,36^. However, a small number of adult escapers survived and exhibited a characteristic rough eye phenotype, overall behaving slightly more strongly than the toolkit attP40 *Ras^G12V^* (Fig. S3A-D). Comparing between the *UAS-Ras^G12X^*toolkit lines, we found that the G12D variant formed pupae at 25°C, but failed to eclose as adults, whereas the G12A, G12C, G12S and G12V variants showed significant lethality (Fig. 3A,B). Viability of the *GMR*>*Ras^G12R^* variant at 25°C was indistinguishable from the *GMR*>*w*^1118^ and *GMR*>*Ras^WT^* controls, whereas all *GMR*>*Ras^G12X^* lines were viable at the slightly milder 21°C (Fig. 3A,B). While *GMR-GAL4* principally drives in the developing visual system, this line has also been shown to express in other tissues^37^, which could explain this lethality.

**Figure 3.**
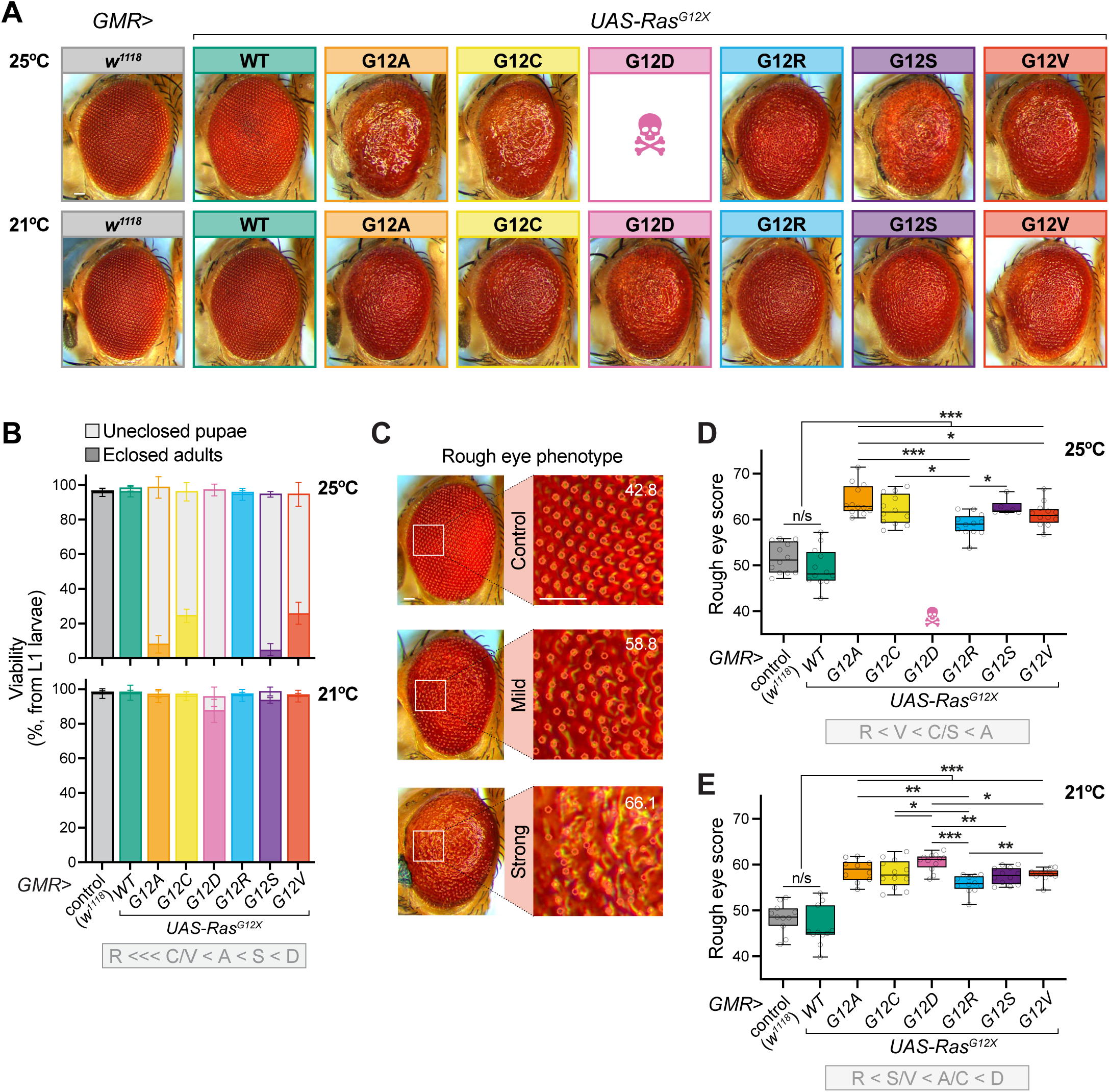
Expression of Ras^G12X^ variants in the developing eye induces differential morphological defects. (**A**) Representative z-stack images of adult female eyes, eclosed following incubation at 25°C or 21°C. The *GMR-GAL4* driver was used to express the *UAS-Ras^G12X^* transgenes in the developing visual system (scale bar: 50 μm). (**B**) Scoring of total viability to pupae and eclosed adults (shaded and solid colour-coding, respectively) at 25°C or 21°C. Data are means ±SD of n=4 vials per condition, each with 50 picked L1 larvae. (**C**) Example outputs of the Flynotyper tool, used to quantify the rough eye phenotype in the *GMR-GAL4*> *UAS-Ras^G12X^*flies (scale bar: 50 μm). The corresponding phenotypic score is displayed as an inset. (**D**,**E**) Automated scoring of eye roughness at 25°C (**D**) and 21°C (**E**) using Flynotyper. Data are boxplots (line at median, interquartile range, min-to-max whiskers) of n=6-12 females per condition, analysed by one-way ANOVA. n/s, p>0.05; *, p<0.05; **, p<0.01; ***, p<0.001.

We imaged the eyes and performed computational analysis using the automated Flynotyper tool^38^ to quantify ‘roughness’ and refine our stratification of phenotypic severity between the Ras^G12X^ variants (Fig. 3C). At 25°C, since *GMR*>*Ras^G12D^* was lethal, the Ras^G12A^ mutant exhibited the most severe rough eye phenotype, followed by the Ras^G12C^ and Ras^G12S^ variants, then Ras^G12V^ (Fig. 3D). At 21°C, *GMR*>*Ras^G12D^* was the most severe, followed by the Ras^G12A^ and Ras^G12C^ variants, then Ras^G12S^ and Ras^G12V^ (Fig. 3E). Ras^G12R^ was the mildest at both 21°C and 25°C (Fig. 3D,E).

### Effect of Ras^G12X^ on development (3): Wing developmental phenotyping

We also characterised our Ras^G12X^ toolkit in the *Drosophila* wing, which is widely used as a model system to study tissue development, growth and patterning^39,40^. The *ptc-GAL4* (*patched*) driver induces Ras^G12X^ expression in a central region of the larval wing imaginal disc, forming a stripe along the anterior-posterior compartment boundary (Fig. 4A). Note that the *ptc*>*Ras^G12V^*(P) combination showed strong developmental lethality (Fig. S4A,B), and therefore was not investigated further. Targeted expression of Ras^WT^ in this region (*ptc*>*Ras^WT^*) did not cause any morphological changes compared to control (*ptc>w*^1118^) wing discs (Fig. 4B,C). In contrast, expression of the various Ras^G12X^ variants (*ptc>Ras^G12X^*) led to developmental (Fig. S4A-C) and morphological defects, including stripe expansion and migration of transformed cells (Fig. 4B,C). Expression of Ras^G12D^ and Ras^G12S^ displayed more severe disorganisation of the *ptc* domain and an aggressive migration of transformed cells (Figs. 4B,C and S4D,E). Intriguingly, the whole wing disc for the Ras^G12R^ variant was significantly smaller in size, despite overall body size of the L3 larvae remaining similar (Fig. S4C,E).

**Figure 4.**
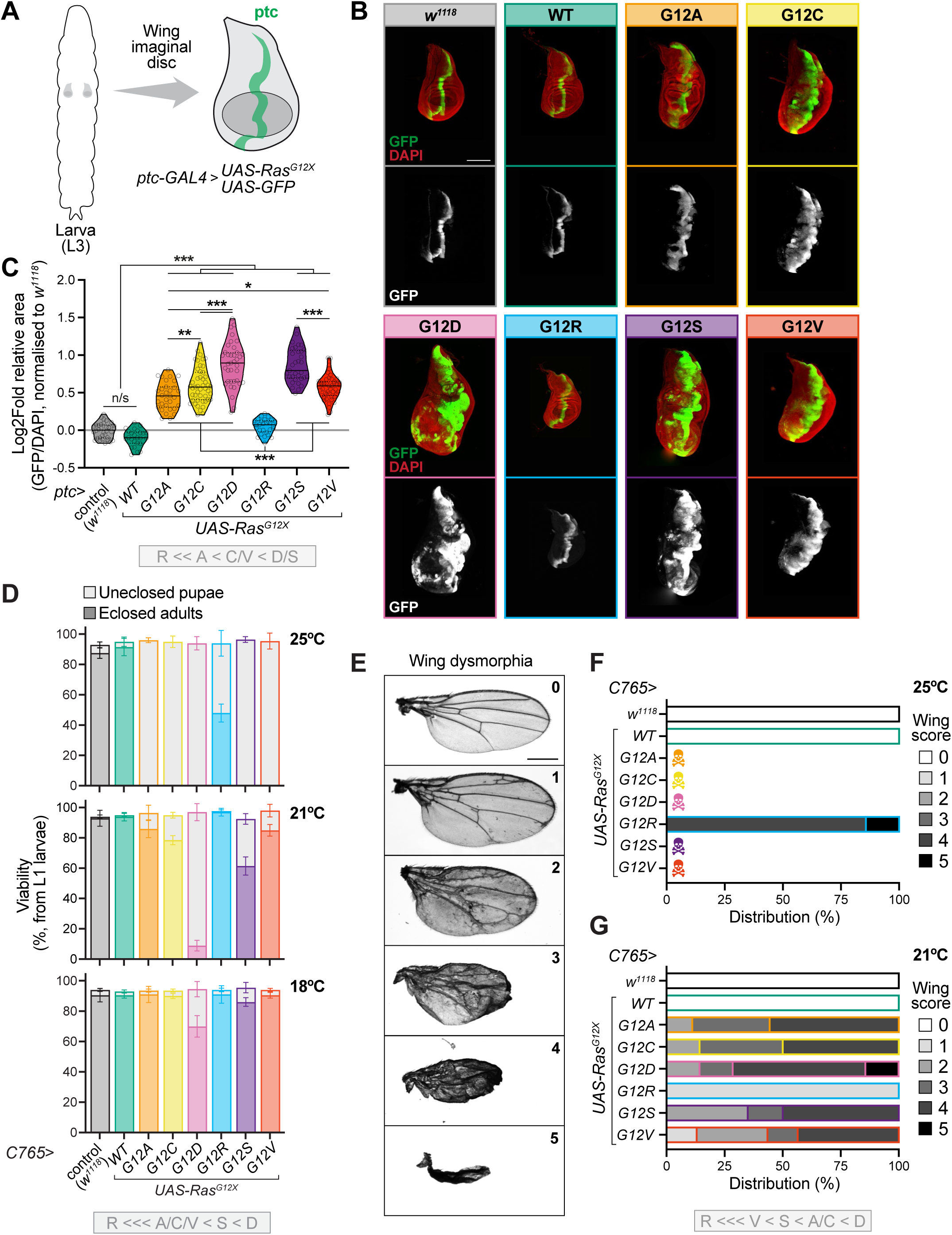
Tissue-specific expression of Ras^G12X^ variants differentially affects wing development. (**A**) The *ptc* enhancer drives the *UAS-Ras^G12X^* transgenes and a *UAS-GFP* reporter in a discrete region of the wing imaginal disc. (**B**) Upper panels: Representative confocal images of DAPI-stained wing imaginal discs (red) dissected from developmental stage-matched L3 larvae, showing the ptc domain labelled by GFP (green). Expression of the *Ras^G12X^* variants leads to distortion and expansion of the ptc stripe (scale bar: 200 μm). Lower panels: Maximum intensity projection of confocal z-stacks showing the GFP channel from above (white). (**C**) Quantification of the ptc domain (GFP), relative to total wing disc area (DAPI), normalised to the *ptc*>*w*^1118^ control (see Fig. S4D,E for the absolute areas). (**D**) The *C765-GAL4* driver expresses in the entire developing wing. Scoring of total viability to pupae and eclosed adults (shaded and solid colour-coding, respectively) at 25°C, 21°C and 18°C. Data are means ±SD of n=4 vials per condition, each with 50 picked L1 larvae. (**E**) Representative images of the scoring system to assess wing dysmorphia, ranging from 0 (wild-type control) to 5 (severe). (**F**,**G**) Quantification of wing defects in *C765*>*Ras^G12X^*females, according to the scale described in **E**. Data for 25°C (**F**) and 21°C (**G**), with n=6-24 flies per condition. n/s, p>0.05; *, p<0.05; **, p<0.01; ***, p<0.001.

We also expressed the *UAS-Ras^G12X^* transgenes using a different wing driver, *C765-GAL4*, across the entire developing wing epithelium. Larvae were cultured over a range of temperatures (18°C, 21°C or 25°C), and assessed for effects on viability (Figs. 4D and S4F) and subsequent defects in adult wing morphology according to a severity scale (Fig. 4E). As expected from the literature, expression of the original *UAS-Ras^G12V^*(P) P-element line caused significant lethality and wing dysmorphia with the *C765-GAL4* driver^41^. The impact on viability was stronger than the toolkit attP40 *Ras^G12V^* line, whereas the wing defects at 21°C were slightly less severe (Fig. S4F,G). Within the toolkit, *C765>Ras^WT^*was indistinguishable from the *C765>w*^1118^ control at all tested temperatures, in terms of viability to adulthood (Fig. 4D). At 25°C, all *C765>Ras^G12X^*variants were lethal during the pupal stage, except *C765>Ras^G12R^*, where ∼50% adults eclosed, but exhibited strong wing deformations (Fig. 4F). At 21°C, G12D exhibited the strongest wing dysmorphia, with severe blistering and crumpling, while G12R was the mildest, with a fully straight wing but still exhibiting venation defects (Fig. 4G).

### Stratification of Ras^G12X^ variants in *Drosophila* tumour contexts (1): Larval Ras/Src co-activated eye disc model

Having explored how the Ras^G12X^ variants differentially affect developmental processes, we next focused on another important physiological impact of Ras: its role in tumour growth and transformation. Therefore, we set out to compare the oncogenic properties of the Ras^G12X^ variants across multiple established tumour models in *Drosophila*. The majority of *Drosophila* studies to-date almost exclusively study the G12V mutation, resulting in a profound knowledge gap of how other Ras^G12X^ variants behave comparatively.

In previous work, we utilised a Ras/Src co-activated tumour model^42,43^, where the original P-element transgene of oncogenic G12V (*UAS-Ras^G12V^*(P)) ^28^ was combined with *csk^Q^*^156^*^Stop^* (null-allele of C-terminal Src-kinase, a negative regulator of Src)^44^, to create GFP-labelled clones specifically in the eye disc under control of the *eyeless* enhancer. As reported^42^, Ras/Src co-activation demonstrated a strong growth response when exposed to a high-sugar diet (HSD) (Fig. S5A-C). In addition to aggressive overgrowth of the primary tumour in the eye epithelia, these *Ras^G12V^*(P)*;csk^−/−^* larvae also displayed ectopic tumours on a HSD (Fig. S5B, arrowheads)^42^. Importantly, this initial study only assessed the G12V variant. Therefore, we combined our *UAS-Ras^G12X^* toolkit with the required *dCsk^Q^*^156^ line, to allow generation of GFP-labelled Ras/Src co-activated eye disc clones for each Ras^G12X^ variant. To assess oncogenic severity, we measured: 1) the degree of primary tumour overgrowth (based on GFP fluorescence), and 2) the occurrence of ectopic tumours on a HSD. Similar to the original P-element *Ras^G12V^*(P)*;csk^−/−^* larvae, our toolkit *Ras^G12V^;csk^−/−^*larvae responded to HSD-treatment with significant tumour enlargement, albeit reaching a comparatively smaller final tumour size (Fig. S5B,C).

We compared tumours generated by combining each of our toolkit Ras^G12X^ variants with Src activation (*Ras^G12X^;csk^-/-^*). Clones with WT Ras and elevated Src activity (*Ras^WT^;csk^-/-^*) were rarely obtained in animals fed either a control diet or HSD (Fig. 5A,B). In larvae fed a control diet, the combination of oncogenic Ras^G12X^ variants with Src co-activation (*Ras^G12X^;csk^-/-^*) resulted in a mild increase in clone size compared to Ras^WT^/Src (Fig. 5A-C). Interestingly, Ras^G12X^/Src clones exhibited variable growth responses to HSD treatment. Specifically, G12A and C12C displayed the most pronounced growth response on the HSD when co-activated with Src (Fig. 5A,B,D). Strikingly, while most oncogenic Ras^G12X^/Src combinations exhibited enhanced expansion of tumours under HSD conditions relative to their counterparts on a control diet, Ras^G12R^/Src-clones instead showed a decreased relative growth response on a HSD (Fig. 5A-D).Next, we assessed the number of GFP^+^ ectopic tumours distant from the eye disc, which can occur upon HSD treatment (Fig. 5A, arrowheads). Using fluorescence microscopy analysis of intact larvae, we quantified the presence and number of these ectopic tumours per larva, and developed a scoring system to classify their size (Fig. S5D-G). Similar to the original P-element Ras^G12V^(P)/Src larvae, our toolkit Ras^G12V^/Src larvae responded to a HSD and displayed GFP^+^ ectopic tumours (Fig. S5B,H,I). Interestingly, while the size of both the primary and ectopic tumours upon HSD-treatment was larger for the P-element *Ras^G12V^*(P)*;csk^−/−^* larvae, the toolkit *Ras^G12V^;csk^−/−^*combination exhibited a higher proportion of larvae with ectopic tumours present (∼90% versus ∼60%), and an increase in larvae with multiple ectopic tumours (Fig. S5B,H,I).

**Figure 5.**
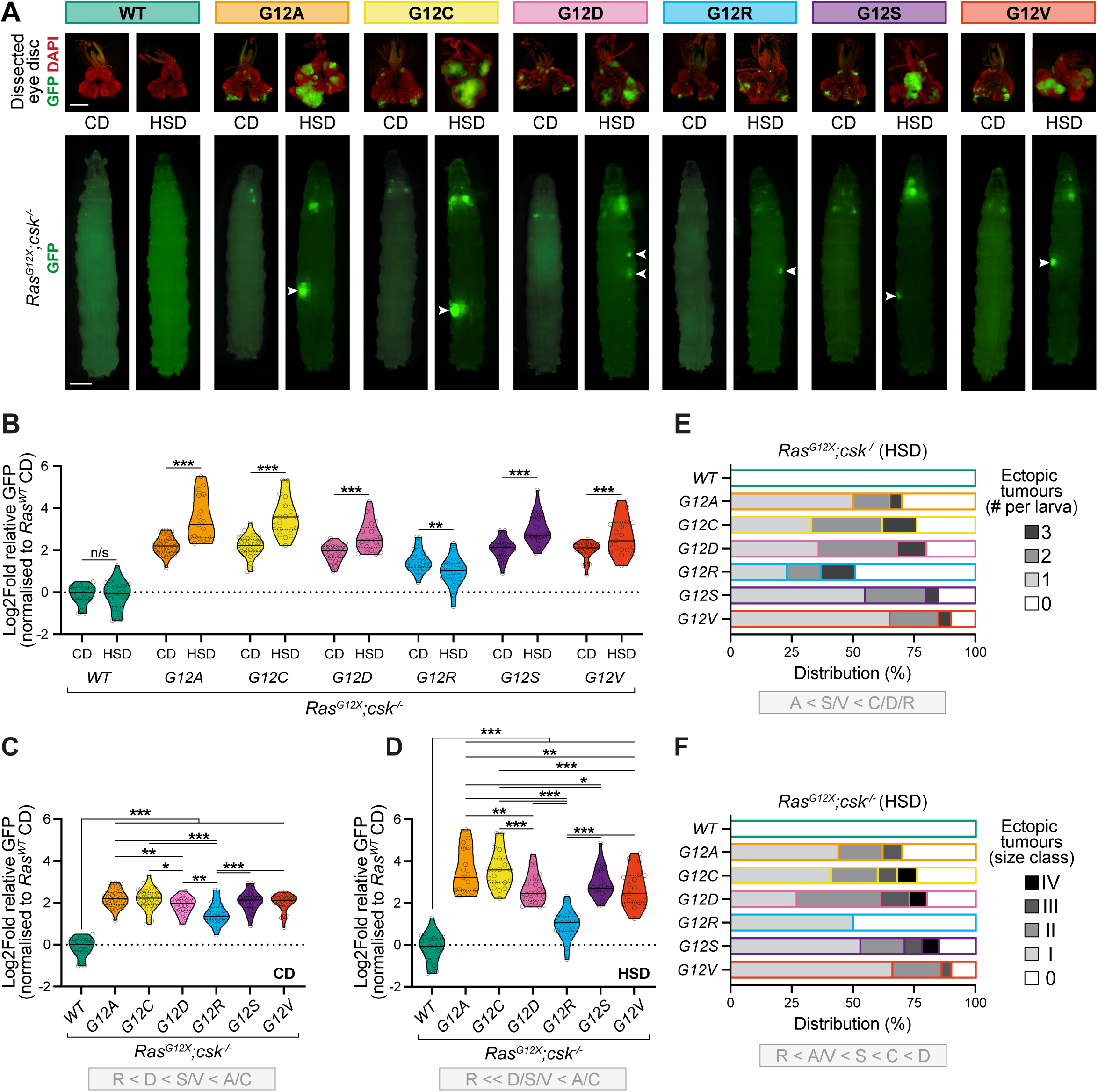
Ras^G12X^/Src co-activated eye disc clones exhibit variable responses to a high-sugar diet (HSD). (**A**) Representative images of developmental stage-matched L3 larvae fed a control diet (CD; day 7 after egg laying) or high-sugar diet (HSD; day 11 after egg laying). The *Ras^G12X^*variants were combined with a *csk^-/-^*(C-terminal Src kinase) mutation to achieve Ras/Src co-activation. Above: Fluorescence microscopy of DAPI-stained dissected eye imaginal discs (red), with transformed tissue labelled by GFP (green) (scale bar: 250 μm). Below: Matched images of whole larvae (scale bar: 500 μm), with ectopic tumours observed in a subset of larvae fed a HSD (arrowheads). (**B**-**D**) Eye disc tumour size for the *Ras^G12X^*/*Src* genotypes, quantified as GFP area relative to the Ras^WT^/Src control, in developmental stage-matched L3 larvae fed on CD or HSD. Data are violin plots (solid line median, dotted line quartiles) of n=17-30 larvae per condition. (**B**) Comparison of CD v HSD, analysed by two-way ANOVA. Statistical comparisons between genotypes within each respective diet, analysed by one-way ANOVA: CD (**C**) and HSD (**D**). (**E**,**F**) Quantification of ectopic tumour number (**E**) and size (**F**) for the different *Ras^G12X^*/Src genotypes. See Fig. S5G for the scoring system. Bar charts for n=20-25 larvae per condition. n/s, p>0.05; **, p<0.01; ***, p<0.001.

The frequency of ectopic tumours in larvae combining our toolkit Ras^G12X^ variants with Src activation (*Ras^G12X^;csk^-/-^*) ranged from 0% for Ras^WT^/Src, ∼50% for Ras^G12R^/Src, up to ∼90% for Ras^G12V^/Src (Fig. 5E). The number of ectopic tumours per larva varied between 1 and 3, with the Ras^G12C^/Src, Ras^G12D^/Src and Ras^G12R^/Src variants showing a higher proportion with multiple foci. For some variants such as Ras^G12R^/Src, we found that the ectopic GFP^+^ clones remained small, while others (e.g. Ras^G12C^/Src and Ras^G12D^/Src) tended to exhibit much larger tumours (Fig. 5F). Note that overall larval body size was broadly unchanged on a HSD across the Ras^G12X^/Src mutants compared to the Ras^WT^/Src control genotype, except Ras^G12R^/Src, which showed a modest decrease, and Ras^G12V^(P)/Src, which was slightly larger (Fig. S5F). Together, these results indicate that different oncogenic Ras^G12X^ variants strongly influence the outcome of Ras/Src-mediated tumour responses to a HSD.

### Stratification of *Ras^G12X^* in *Drosophila* tumour contexts (2): Proliferation of ISCs in the adult midgut

To complement the above larval tumour model, we next studied behaviour of the Ras^G12X^ variants in two established adult tumour models: for the midgut (=*Drosophila* small intestine) and the hindgut (=*Drosophila* colon)^45^. Both these systems are well recognised in the field (e.g. ^29,30,46,47^). However again, while there is abundant data for G12V, the other G12X mutations remain uncharacterised.

First, we studied how the Ras^G12X^ variants impact the proliferation of midgut intestinal progenitor cells, using the ISC (intestinal stem cell)/EB (enteroblast) driver *esg-GAL4* (*escargot*), and scoring for phospho-histone-H3^+^ cells (pH3^+^), a marker of mitosis, after 7 days of induction (Fig. 6A,B). A ubiquitous temperature-sensitive repressor (*tub-GAL80^ts^*) prevents expression during development^48^. As expected from the literature^29^, the original P-element *esg^ts^*>*Ras^G12V^*(P) caused a strong proliferation phenotype, behaving similarly to the toolkit attp40 *Ras^G12V^* line in this adult model (Fig. S6A-C). Interestingly, for this paradigm, our data indicate that Ras^G12R^ (typically amongst the mildest variants) and Ras^G12V^ were the most severe in terms of proliferation, based on pH3^+^ midgut counts (Fig. 6C,D). Survival-wise, Ras^G12V^ was the shortest-lived variant, clustering with Ras^G12D^, the latter exhibited a more moderate pH3^+^ scoring. Conversely, Ras^G12R^ was the longest-lived variant, despite a strong pH3^+^ phenotype (Fig. 6C-E). Therefore, in the context of this midgut tumour model, the severity of proliferation and subsequent survival can be partially uncoupled.

**Figure 6.**
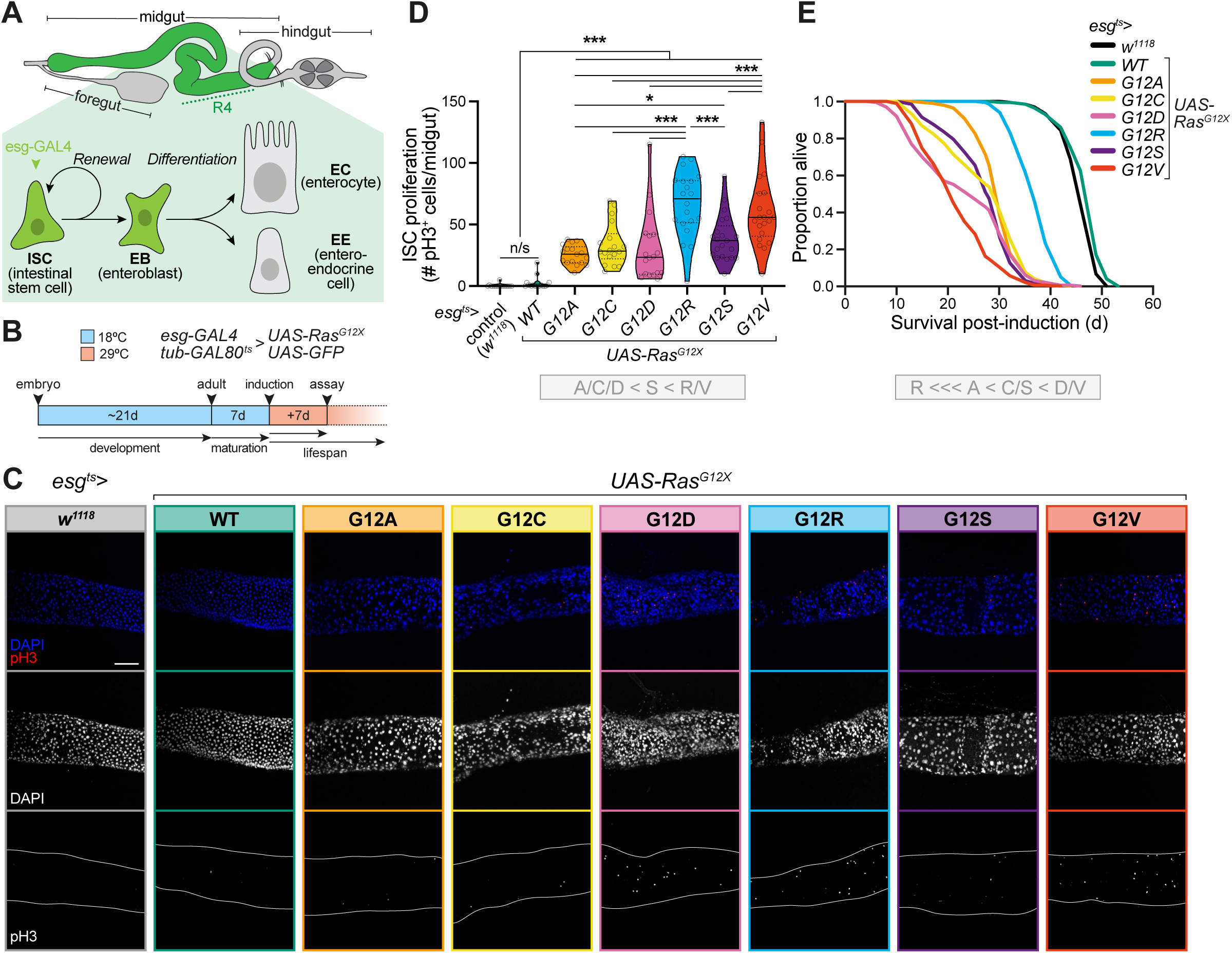
Impact of Ras^G12X^ variants on proliferation of midgut ISCs. (**A**) Diagram of the adult midgut tumour model. The *esg-GAL4* driver expresses the *UAS-Ras^G12X^* transgenes and a *UAS-GFP* reporter in the self-renewing midgut intestinal stem cells (ISCs) and their undifferentiated transient daughters, enteroblasts (EB). Combination with the temperature-sensitive *tub-GAL80^ts^* system bypasses developmental effects. EC, enterocyte, EE, entero-endocrine cell. (**B**) Scheme of the experimental set-up. Flies were assayed after 7 days of induction at 29°C. (**C**) Representative confocal images of pH3^+^ cells (red), indicative of proliferation (mitosis), and DAPI staining of all nuclei (blue) for *esg^ts^*>*UAS-Ras^G12X^* females. The R4 region of the posterior midgut is shown (scale bar: 100 µm). (**D**) Scoring of pH3^+^ cells across the entire midgut for the Ras^G12X^ variants. Data are violin plots (solid line median, dotted line quartiles) of n=16-21 females per genotype, analysed by one-way ANOVA. (**E**) Survival curves post-tumour induction for *esg^ts^*>*UAS-Ras^G12X^*. Lifespans were performed with n=∼200 females per condition (see Table S6 for full details of n numbers and Log-Rank p values). n/s, p>0.05; *, p<0.05; ***, p<0.001.

### Stratification of Ras^G12X^ in variants *Drosophila* tumour contexts (3): Adult hindgut model of colorectal cancer

Finally, we explored an established *Drosophila* model of colorectal cancer, which has previously been used to investigate how tumours respond to bacterial infection and metabolites^30,49^, and how complex mutational profiles determine the response and resistance to targeted therapies^47^. Here, the *UAS-Ras^G12X^*variants were expressed under control of the hindgut-specific driver *byn-GAL4*, combined with *tub-GAL80^ts^* (*byn ^ts^*) to restrict expression during adulthood. Newly eclosed adult flies were maintained at 18°C for 7 days to allow maturation, then transgene expression was induced by switching to 29°C. Additionally, a GFP-reporter was included for visualisation of both primary tumour growth and secondary tumour invasiveness. Phenotypes were assessed following induction for 14 days (Fig. 7A,B).

**Figure 7.**
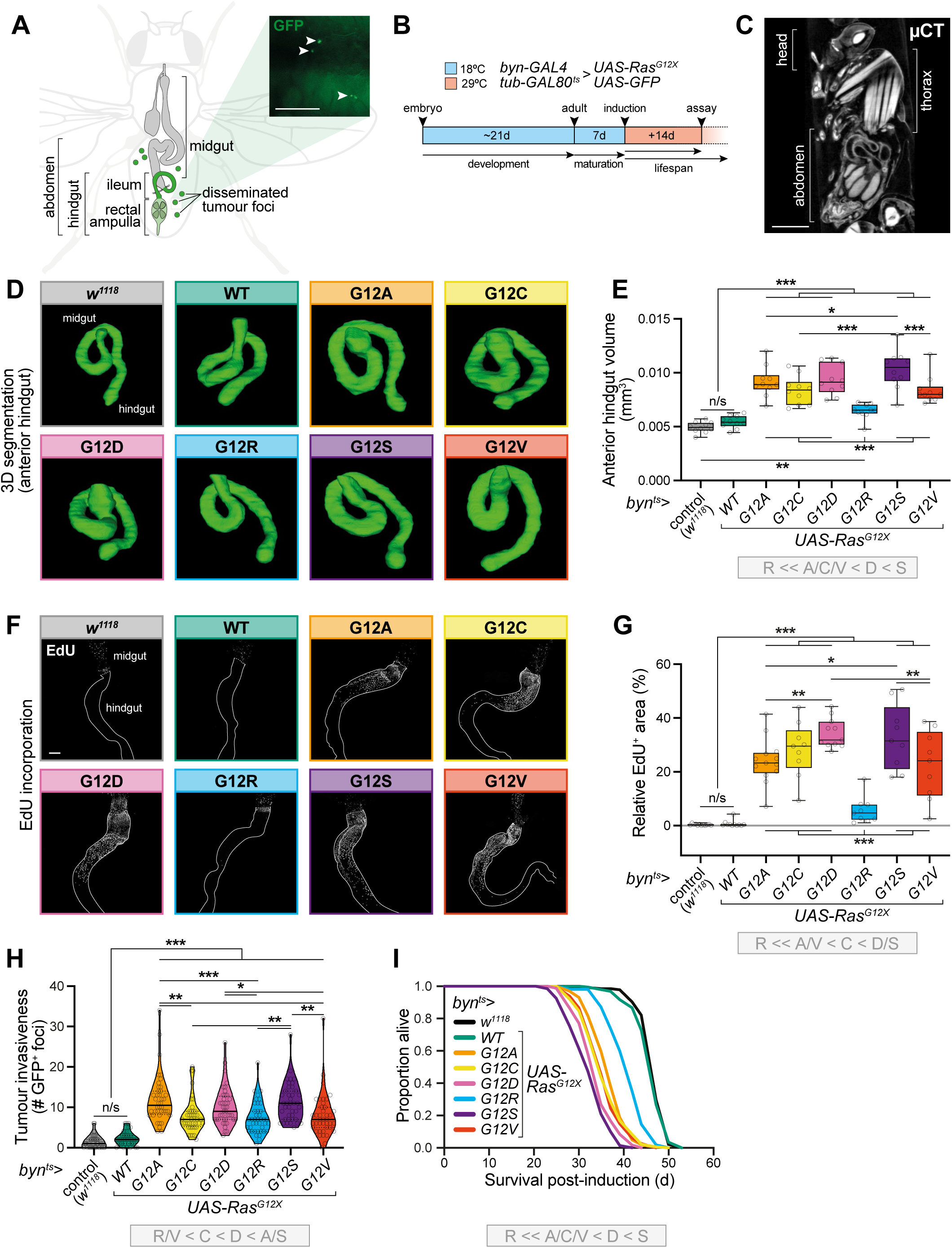
Impact of Ras^G12X^ variants in an adult colon cancer model. (**A**,**B**) Scheme of the experimental system. (**A**) The hindgut-specific driver, *byn-GAL4*, drives the *UAS-Ras^G12X^* transgenes, combined with the temperature-sensitive *tub-GAL80^ts^* system to prevent expression during development. Co-expression with a *UAS-GFP* reporter labels the primary tumour and fluorescent foci that disseminate from the hindgut into the abdomen upon Ras activation (inset: example fluorescence microscopy image of GFP^+^ foci, scale bar: 100 µm). (**B**) Experimental timeline. (**C**-**E**) Micro-computed tomography (µCT) imaging of adult hindguts *in situ*. (**C**) Example 2D cross-section of an adult female (scale bar: 500 µM). (**D**) Representative 3D reconstruction of an anterior hindgut (defined as the pylorus valve and ileum, excluding the rectal ampulla) for each *byn^ts^*>*UAS-Ras^G12X^* variant. (**E**) Quantification of anterior hindgut volume by µCT. Data are boxplots (line at median, interquartile range, min-to-max whiskers) for n=10 females per genotype, analysed by one-way ANOVA. (**F**,**G**) Assessment of cell proliferation based on EdU incorporation. (**F**) Typical confocal images of hindguts, outlined by solid lines, and the midgut/hindgut junction indicated by a dotted line (scale bar: 100 µm). EdU^+^ cells are only present in the midgut of control flies (*byn^ts^*>*w*^1118^ and *byn^ts^*>*UAS-Ras^WT^*), but expand into the hindguts of *byn^ts^*>*UAS-Ras^G12X^*variants. (**G**) Quantification of the EdU^+^ area, relative to the total anterior hindgut. Data are boxplots (line at median, interquartile range, min-to-max whiskers) for n=9-13 flies per genotype, analysed by one-way ANOVA. See Fig. S7C for the corresponding DAPI staining. (**H**) Dissemination assay data comparing the *Ras^G12X^* variants. Data are violin plots (solid line median, dotted line quartiles) of n=40 females per genotype, analysed by a Kruskal-Wallis test. (**I**) Survival curves of *byn^ts^*>*UAS-Ras^G12X^*flies post-tumour induction from d7 of adulthood. Lifespans were performed with n=∼140-200 females per condition (see Table S6 for full details of n numbers and Log-Rank p values). The µCT imaging (**D**,**E**), EdU staining (**F**,**G**) and dissemination data (**H**) were performed on *byn^ts^*>*UAS-Ras^G12X^*females induced from d7 of adulthood for 14 days at 29°C. n/s, p>0.05; *, p<0.05; **, p<0.01; ***, p<0.001.

Firstly, to quantify the size of the primary hindgut tumour, we scanned the flies by micro-computed tomography (µCT), a technique that uses X-rays to create high-resolution 3D imaging of internal organs (Fig. 7C)^50–52^. Based on µCT scan segmentation, the original P-element *Ras^G12V^*(P) line exhibited an increase in volume of the anterior hindgut (defined as the pylorus valve and ileum, excluding the rectal ampulla) to a greater extent than the toolkit *Ras^G12V^* line (Fig. S7A,B). Similarly, the anterior hindguts of all toolkit *byn^ts^*>*Ras^G12X^* variants were significantly enlarged upon oncogenic Ras^G12X^ activation, with the Ras^G12S^ and Ras^G12D^ variants showing the strongest increase in volume, reaching ∼0.1 mm^3^ compared to ∼0.005 mm^3^ for the *byn^ts^*>*w*^1118^ and *byn^ts^*>*Ras^WT^*controls (Fig. 7D,E). In contrast, the Ras^G12R^ variant produced the mildest phenotype, only reaching an average anterior hindgut volume of ∼0.007 mm^3^. To expand on these findings, we also assessed proliferation by performing EdU incorporation analysis, which has been published for the P-element *Ras^G12V^*(P) line (with equivalent BrdU)^47^, but is unknown for the remaining Ras^G12X^ variants. Following oncogenic Ras^G12X^ activation in the hindgut, EdU^+^ replicating cells, which are normally restricted to the midgut, were detected in the hindgut (Figs. 7F,G and S7C-E). Here, flies were induced for a total of 14 days, and fed EdU in their diet for the last 7 days prior to quantification. Consistent with our µCT data, the Ras^G12S^ and Ras^G12D^ variants showed the strongest proliferative phenotype based on EdU^+^ staining in the hindgut, whereas Ras^G12R^ was substantially the weakest.

Next, to assess tumour invasiveness, flies were dissected and scored under the fluorescence microscope for GFP^+^ foci, representing hindgut cells that exhibit an invasive behaviour and disseminate throughout the abdominal cavity (Fig. 7A inset, arrowheads)^30,47^. Flies expressing Ras^WT^ behaved as the driver-only control, with baseline levels of foci. Across the Ras^G12X^ variants, Ras^G12S^ exhibited the strongest dissemination phenotype along with Ras^G12A^, while Ras^G12R^ was the least invasive along with Ras^G12V^ (Fig. 7H). Overall, the pattern of tumour dissemination shows some differences compared to the primary hindgut growth and proliferation data, indicating that the relative stratification of the Ras^G12X^ variants varies depending on the tumour phenotype assessed.

Finally, we also performed parallel lifespan assays of the *byn^ts^*>*Ras^G12X^*flies to monitor survival following tumour induction (Fig. 7I). Consistent with the overarching trends in this model, the Ras^G12S^ and Ras^G12R^ variants were the shortest- and longest-lived, respectively. However, the relationship between tumour severity and survival is not straight-forward, since the P-element *Ras^G12V^*(P) line exhibited stronger hindgut proliferation and dissemination than our toolkit *Ras^G12V^* line, but both decreased lifespan to similar extents (Fig. S7A-G).

## Discussion

Human KRAS^G12X^ mutations are differentially distributed across tumour tissue types^5,7^, yet the biological and pharmacological implications of this variation remain underexplored, since experimental models tend to consider only a single or limited subset of KRAS^G12X^ variants. Our novel *Drosophila* toolkit provides unprecedented insight into the behaviour of different Ras^G12X^ variants *in vivo*, enabling systematic decoding of their respective physiological properties. Importantly, *Drosophila* exhibits strong evolutionary conservation with mammals, especially relating to key metabolic and signalling pathways. With its powerful genetic tools, cost-effective and ethical benefits, combined with a capacity for high-throughput phenotypic screening, *Drosophila* is an ideal system to interrogate the behaviour of Ras^G12X^ variants *in vivo*.

For all experimental contexts and read-outs, the *driver>Ras^WT^*condition behaved identically to the endogenous control, *driver*>*w*^1118^. Therefore, overexpression of WT Ras does not disrupt normal physiological development and, moreover, is evidently not sufficient to induce the proliferative and tumour phenotypes - an activating oncogenic G12X mutation is required.

Our study clearly highlights significant tissue-dependent effects. While G12R is typically the weakest variant and G12D often the strongest, the full stratification of Ras^G12X^ variants differs considerably between *in vivo* contexts, as illustrated by the heatmap summary (Fig. 8). All the G12X mutations included in the toolkit exhibit loss of GAP-stimulated GTP hydrolysis, effectively locking Ras in a constitutively active state (Fig. 1A). WT Ras also has the ability to self-inactivate (i.e. independently of GAP). According to the literature, this intrinsic GTP hydrolysis is severely abolished in the G12A, G12R and G12V variants, moderately impaired in G12D, but comparatively preserved in G12C (note that G12S was not included in this study)^8^, therefore suggesting that G12C can cycle between GDP and GTP more readily than other G12X mutants. Another *in vitro* study comparing intrinsic GTP hydrolysis across a panel of KRAS mutations relative to Ras^WT^ reported that this rate was strongly impaired in the G12R and G12V variants (∼20- and ∼10-fold decrease, respectively), while the effect was less pronounced for G12A, G12C and G12D (≤3-fold)^53^.

**Figure 8.**
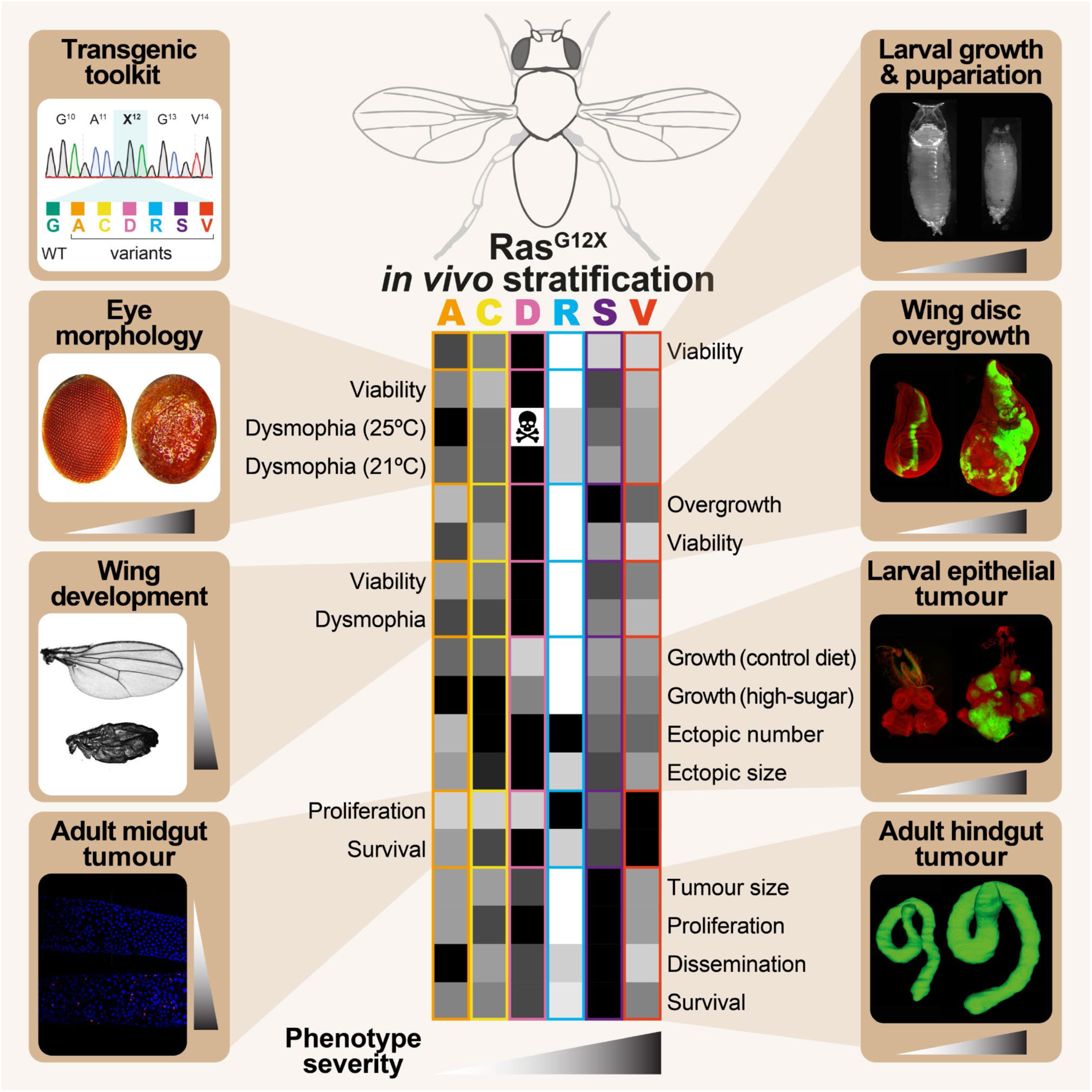
Summary heatmap of the Ras^G12X^ variants. Graphical heatmap summarising the context-dependent functional hierarchies of Ras^G12X^ variants according to their relative phenotypic strength across developmental and cancer contexts.

Codon 12 is within the highly conserved phosphate-binding P-loop domain of KRAS (Fig. S1A). Since glycine is the smallest and simplest amino acid, any G12X substitutions will cause steric hindrance within the pocket where GTP hydrolysis occurs. For instance, G12V (valine) has a bulky hydrophobic side-chain, while G12R (arginine) introduces a disruptive positively charged side-chain, which protrudes directly into the pharmacologically important ‘Switch II’ pocket^54^. Interestingly, due to this structural distortion, G12R displays impaired activation of PI3Kα signalling in pancreatic cancer (PDAC) cell lines^9^, as well as blunted ERK signalling, resulting in a weaker transcriptional growth signal compared to more aggressive mutations such as G12D^55^. To adapt to these downstream pathway defects, G12R-mutated cells are reported to rely heavily on autophagy^9^. Interestingly, PDAC patients with the G12R mutation are associated with better survival than G12V and G12D^56^, consistent with our data showing that G12R is phenotypically the weakest variant.

Biochemical profiling of the G12X variants for their interaction with RAF, a major downstream effector kinase, reveals that G12D, G12R and G12V exhibit extremely poor RAF binding affinity (∼5-7-fold weaker), while G12A was relatively preserved (∼2-fold weaker) and G12C was essentially unaffected (note that G12S was not included in this study)^8^. Indeed, G12A (alanine) has a small non-polar hydrophobic side-chain, enabling Ras to retain flexible open conformations. Hence, G12A is predicted to have strong RAF pathway engagement (i.e. MAPK/ERK signalling). An epidemiological investigation of colorectal cancer revealed that patients carrying the G12A mutation exhibited early recurrence and poor survival outcomes^57^. In another study of patients with metastatic colorectal cancer, the G12S variant was associated with the shortest overall survival^58^. These clinical findings are consistent with our hindgut tumour data (Fig. 7), where the G12A and G12S exhibited strong dissemination phenotypes.

G12D (aspartic acid) introduces a negatively charged side-chain, altering its ‘Switch II’ conformation^59^ and impacting its interaction with downstream effectors. G12D is reported to exhibit strong PI3K/AKT signalling, with less favouring of RAF pathways than G12A and G12C. Pancreatic cancer (PDAC) patients carrying G12D mutations had worse overall survival compared to those with G12R^56^. Additionally, G12D is reported to promote lateral segregation and enhanced dimerisation of KRAS, which impacts recruitment of downstream effectors^60^. In total, more than 50 proteins have been identified as Ras effectors in humans, which exhibit variable expression across tissue types^61,62^. This diversity highlights the potential complexity of downstream signalling processes. All these interactors bind via the same effector domain in Ras, but with a range of affinities^9,61,62^.

Consistent with our findings with the *Drosophila* hindgut (Fig. 7), a study comparing G12C, G12D and G12R in a colorectal cancer mouse model reported that while the G12C and G12D variants induced widespread hyperplasia, the G12R resembled WT tissue^15^. When induced in the mouse pancreas, the G12R variant also had the lowest tumour-initiating and disease-progression capacity compared with the G12C and G12D variants^15^. Another study comparing organoids derived from G12D and G12R variant-induced pancreatic tissue demonstrated that the G12R variant exhibited reduced migratory behaviour and improved overall survival when transplanted orthotopically into mice^63^.

While G12R was consistently the phenotypically weakest variant across the majority of *in vivo* models considered with our toolkit, we interestingly observed an unexpectedly strong pH3^+^ signal with G12R in the midgut tumour model (Fig. 6). A recent study used an analogous *Drosophila* adult midgut cancer model to compare the Ras^G12V^ variant with Ras^Q61H^, another mutation occurring in human tumours affecting a key catalytic site located within the ‘Switch II’ region (Fig. S1A)^64^. While induction of the Ras^Q61H^ variant in ISCs resulted in a stronger, hyperproliferative phenotype, these rapidly expanding cells were eventually eliminated from the midgut. Therefore, longer-term, the comparatively weaker Ras^G12V^ was actually more deleterious than Ras^Q61H^, causing severe intestinal barrier dysfunction^64^. This observation could explain why the typically weaker G12R variant exhibited higher pH3^+^ counts. As our experiments assessed tumour proliferation at 7 days post-induction, future studies examining earlier and later time points would help define the temporal dynamics of G12R-driven proliferation and determine how these compare with the other Ras^G12X^ variants.

Our transgenic constructs include Flag and HA tags, therefore future research can include proteomic analysis of Ras^G12X^ interactors, in order to explore the downstream molecular effectors underlying our phenotypic stratification. Findings can subsequently be translated and explored in a more informed and targeted approach with mammalian organoids or mice.

Longer-term, this Ras^G12X^ toolkit can be further combined with additional oncogenic and/or tumour suppressor mutations (e.g. APC, p53, PTEN), building on our proof-of-principle with Ras/Src co-activation (Fig. 5). Indeed, the *Drosophila* system allows researchers to achieve complex multigenic combinations at an ambitious scale^32,47,65–67^, which would not be practicable or economically viable in equivalent mammalian models.

Ultimately, dissecting the functional differences between Ras^G12X^ variants *in vivo* will improve our fundamental understanding of cancer biology, and will contribute towards developing improved therapeutic strategies targeting patients with specific KRAS^G12X^ mutations. The emergence of variant-specific therapeutics highlights the importance of unravelling the complexity of the Ras^G12X^ code. The *Drosophila* system is already used to generate patient-specific avatars in order to screen for effective drugs (e.g. ^32,65,68^), and this Ras^G12X^ toolkit can further guide translational endeavours at personalised medicine for tumours carrying defined KRAS^G12X^ mutations.

## Methods

| **Experimental models: *Drosophila* lines** | **Source** | **Identifier/Reference** |
|---|---|---|
| *w<sup>1118</sup>* | Linda Partridge | N/A |
| *da-GAL4* (Chr 3, backcrossed into *w<sup>1118</sup>*) |  | Wodarz *et al.* (1995)<sup>69</sup> |
| *da-GS* (Chr 2, backcrossed into *w<sup>1118</sup>*) |  | Tricoire *et al.* (2009)<sup>70</sup> |
| *GMR-GAL4* (Chr 3, backcrossed into *w<sup>1118</sup>*) | Bloomington #8605 | RRID:BDSC_8605 |
| *C765-GAL4* (Chr 3, backcrossed into *w<sup>1118</sup>*) | Bloomington #36523 | RRID:BDSC_36523 |
| *ptc-GAL4* (Chr 2, backcrossed into *w<sup>1118</sup>*) | Bloomington #2017 | RRID:BDSC_2017 |
| *UAS-dcr2;; byn-GAL4,tub-GAL80<sup>ts</sup>,UAS-GFP/TM6b* | Erdem Bangi & Ross Cagan | Bangi *et al.* (2012)<sup>30</sup> |
| *w\*; esg-GAL4<sup>NP7397</sup>,UAS-GFP, tub-GAL80<sup>ts</sup>/CyO* | Bruce Edgar | Jiang & Edgar (2009)<sup>71</sup> |
| *dCsk<sup>Q156Stop</sup>* (Chr 3) | Alana O'Reilly & Michael Simon | O'Reilly *et al*. (2006)<sup>44</sup> |
| *ey(3.5)-FLP1; act > y<sup>+</sup> > GAL4,UAS-GFP; FRT82B, tub-GAL80* | Ross Cagan | Hirabayashi *et al.* (2013)<sup>42</sup> |
| *UAS-Ras<sup>WT</sup>* (*UAS-Flag-His-Ras<sup>WT</sup>*) (Chr 2 attP40) (backcrossed into *w<sup>1118</sup>*) | This study | N/A |
| *UAS-Ras<sup>G12A</sup>* (*UAS-Flag-His-Ras<sup>G12A</sup>*) (Chr 2, attP40) (backcrossed into *w<sup>1118</sup>*) | This study | N/A |
| *UAS-Ras<sup>G12C</sup>* (*UAS-Flag-His-Ras<sup>G12C</sup>*) (Chr 2, attP40) (backcrossed into *w<sup>1118</sup>*) | This study | N/A |
| *UAS-Ras<sup>G12D</sup>* (*UAS-Flag-His-Ras<sup>G12D</sup>*) (Chr 2, attP40) (backcrossed into *w<sup>1118</sup>*) | This study | N/A |
| *UAS-Ras<sup>G12R</sup>* (*UAS-Flag-His-Ras<sup>G12R</sup>*) (Chr 2, attP40) (backcrossed into *w<sup>1118</sup>*) | This study | N/A |
| *UAS-Ras<sup>G12S</sup>* (*UAS-Flag-His-Ras<sup>G12S</sup>*) (Chr 2, attP40) (backcrossed into *w<sup>1118</sup>*) | This study | N/A |
| *UAS-Ras<sup>G12V</sup>* (*UAS-Flag-His-Ras<sup>G12V</sup>*) (Chr 2, attP40) (backcrossed into *w<sup>1118</sup>*) | This study | N/A |
| *UAS-Ras<sup>G12V(P)</sup>* (*=UAS-Ras1<sup>V12</sup>*) (Chr 2, P-element) (backcrossed into *w<sup>1118</sup>*) | Gerald Rubin | Karim & Rubin (1998)<sup>28</sup> |

### Fly Strains

The *Drosophila* lines used in this study are detailed above. The *UAS-Ras^G12X^* transgenic toolkit was generated as follows. The Flag-His6-Ras-WT construct was synthesised by GenScript (https://www.genscript.com) and cloned into the pUASTattB vector using EcoRI/NotI restriction sites. The series of G12X variants (G12A, G12C, G12D, G12R, G12S and G12V) was subsequently generated from the WT by targeted mutagenesis, accounting for *Drosophila* codon preference (see Table S5 for all sequences). Plasmids were injected by BestGene (https://www.thebestgene.com) into embryos of the genotype *y^1^w*^67c23^*;P{CaryP}attP40*, allowing phiC31 integrase-mediated targeted insertion into the attP40 site on chromosome 2. Transformants were selected and balanced with CyO. All lines were backcrossed into the *w*^1118^ background for 6 generation, before finally establishing as homozygote stocks. Presence of the correct point mutations was confirmed by sequencing.

### Fly Maintenance

Experiments were performed with the *w*^1118^ strain of *Drosophila melanogaster* as the control background. UAS and driver lines were typically backcrossed into the *w*^1118^ background for ∼6 generations. Adult assays were mostly performed on mated females, unless otherwise stated. Stocks were maintained long-term at 18°C. Experiments were performed at 18°C, 21°C, 25°C or 29°C, as indicated, in either incubators or controlled temperature rooms on a 12 h light:12 h dark cycle, with 65% humidity.

### Fly Media

For the majority of experiments, flies were raised and maintained on standard sugar-yeast-agar (SYA) medium consisting of: 5% w/v sucrose (granulated sugar), 10% w/v Brewer’s yeast (MP Biomedicals, 903312), 1.5% w/v agar (Sigma, A7002), supplemented with nipagin (p-hydroxy-benzoic acid methyl ester; Sigma, H5501; 30 mL/L of 10% w/v nipagin in 95% EtOH) and propionic acid (0.3% v/v; Sigma, P1386) as preservatives, added once the food had cooled down to <60°C^72^. For dietary drug treatments, SYA was supplemented with 200 µM RU (RU486/mifepristone; Sigma, M8046; from a 100 mM stock in 100% EtOH), or 5 mg/mL EdU (5-ethynyl-2’-deoxyuridine; APExBIO, B8337).

For wing disc experiments (ptc-GAL4), flies were cultured on standard cornmeal-containing medium containing: 6.65% w/v cornmeal (organic corn maize, Doves Farm), 7.15% w/v (glucose; Sigma, G8270), 5% w/v autolysed yeast powder (Kerry, #20050488), 0.66% w/v agar (No 3; Oxoid, LP0013), supplemented with 2.2% v/v nipagin (Sigma, H3647) and 3.4 mL/L of propionic acid (Sigma, P1386)^52^.

For the Ras/Src co-activation experiments, flies were cultured on a modified Bloomington semi-defined medium (https://bdsc.indiana.edu/information/recipes/germanfood.html), with sucrose as the only purified sugar source, as previously described^42,73^. The control diet (CD) and high-sugar diet (HSD) contained 0.15 M and 1.0 M sucrose, respectively. Ingredients were sourced as follows: agar (Fisher Scientific, BP2641-1), Brewer’s yeast (MP Biomedicals, 903312), yeast extract (Sigma, 70161), peptone (Sigma, 82303), sucrose (Fisher Scientific, S/8560/63), magnesium sulfate hexahydrate (Fluka, 00627), calcium chloride dihydrate (Sigma, 223506), propionic acid (Sigma, P1386), nipagin (Sigma, H5501). HSD feeding led to a developmental delay in reaching the L3 larval stage when cultured at 25°C. All experiments were performed on stage-matched late L3 larvae at day 7 after egg laying (CD) or day 11 after egg laying (HSD).

### Experimental Flies

For most experiments, crosses (driver virgins x *w*^1118^ control or UAS males) were set up in cages with apple juice agar bases and a drop of fresh yeast paste to encourage egg laying. Bases were changed regularly (∼once per day). For developmental assays, L1 larvae were picked from apple juice agar plates and transferred to vials using a mounted needle to ensure constant density as follows: n=50 per narrow vial (25 mm diameter) of SYA medium (*da-GAL4*, *GMR-GAL4*, *C765-GAL4*), or n=100 per wide vial (28.5 mm diameter) of cornmeal medium (*ptc-GAL4*). Animals were incubated at the appropriate temperature, then scored for pupariation and eclosion.

For adult experiments, eggs were collected over a <12 h period to ensure a synchronous population and reared at constant density by seeding ∼250 eggs per SYA bottle (16 µL of egg suspension using a wide-bore tip)^72^. For experiments at 25°C, eclosing adults (day 0) of a defined age were kept as a mixed population for ∼2 days on standard SYA to allow maturation and mating, then separated into females and males under mild CO2 anaesthesia, and maintained as separate sexes in vials from then onwards (typically 15 or 20 flies per vial). For experiments at 18°C, flies were kept as a mixed population for ∼3-4 days before sorting. Flies were transferred to fresh food every ∼2-3 days (25°C and 29°C) or ∼3-5 days (18°C) without anaesthesia.

### Survival assays

For survival assays, flies were generated and set up as above, typically with n=20 females per narrow vial and a total of n=100-200 flies per condition. Flies were incubated at the appropriate temperature (29°C for experiments with the inducible *tub-GAL80^ts^* system), and transferred to fresh food without anaesthesia every ∼2-3 days, with deaths and censors recorded.

### Western blotting

Control (*da-GS*>*w*^1118^) or experimental (*da-GS*>*UAS*-*Ras^G12X^*) flies were reared at 25°C on standard SYA medium (–RU) until d7 of adulthood. Ubiquitous Ras^G12X^ expression was induced by transferring flies to SYA supplemented with RU (+RU, 200 µM) for 48 h. Following induction, flies were snap frozen on dry ice and stored at –70°C until required.

Frozen females (x10 adults per sample) were homogenised directly into in 200 µL of 2x Laemmli Sample Buffer (BioRad, #1610737) supplemented with 5% (v/v) β-mercaptoethanol (Sigma, 63689) using a pestle and motor. The samples were heated for 5 min at 95°C, and centrifuged for 2 min at 10,000 rpm, RT. Subsequently, 5 μL of homogenate per condition was resolved by SDS-PAGE using a Criterion TGX Stain-Free precast gel (Any kD; Bio-Rad, #5678123) and transferred onto a 0.2 μm PVDF membrane (Trans-Blot Turbo Midi Format; Bio-Rad).

Following transfer, the membrane was blocked with 5% milk prepared in Tris-buffer saline supplemented with 0.05% Tween20 (TBS-T) for 1 h at room temperature (RT). The membrane was then incubated overnight at 4°C with mouse anti-Pan-Ras primary antibody (1:500; Sigma, OP40) diluted in 5% BSA in TBS-T. Following the overnight incubation, the membrane was washed with TBS-T and incubated with anti-mouse secondary antibody (1:1,000; Cell Signaling Technology, #7076) diluted in 5% milk in TBS-T for 1 h at RT. Subsequently, the membrane was washed, and specific protein bands were visualised using the SuperSignal West Femto chemiluminescent substrate (Thermo Scientific), and imaged on a ChemiDoc Touch Imaging System (Bio-Rad).

### Light Microscopy Imaging

#### Pupal size

Pupae were transferred to a glass microscope slide and imaged on a Leica M165FC microscope at 1.25x magnification using LAS X software (Leica). Length was measured in FIJI.

### Adult eye imaging and analysis

For the adult eye phenotyping (*GMR-GAL4* driver), previously frozen females (∼d1 post-eclosion) were mounted laterally using a strip of petroleum jelly (Vaseline) on a glass microscope slide, with the left eye facing upwards for imaging. Z-stack images were acquired using a motorised Leica M165 FC stereo microscope equipped with a Leica K3 digital camera with a Z focus drive, using a 12x objective. Z-stacks were initially processed using LAS X software (Leica) to generate multi-focus extended-depth-of-field images, and subsequently analysed with the Flynotyper tool^38^ to generate rough eye phenotypic scores (see Fig. 3C).

#### Adult wing imaging and scoring

For the adult wing phenotyping (*C765-GAL4* driver), the right wing was removed with fine forceps from previously frozen females (∼d1 post-eclosion) and mounted onto a glass microscope slide in isopropanol. Wings were imaged using a Leica M165 FC stereo microscope equipped with a Leica K3 digital camera with a Z focus drive, using a 12x objective. Morphology was classified according to a dysmorphia scoring system (see Fig. 4E).

### Immunohistochemistry and analysis of imaginal discs

Stage-matched L3 larval imaginal discs were dissected in 1X PBS (Gibco, 10010-015), fixed in 4% PFA (Thermo Scientific, 28908) for 30 min on ice, and washed x3 times in 1X PBS. Eye discs were incubated in Vectashield with DAPI (Vector Laboratories, H-1200) for 30 min to allow DAPI penetration prior to final mounting. Wing discs were incubated with DAPI (1:1,000; Invitrogen, D1306) in 1X PBS for 2 h and then mounted in Vectashield (Vector Laboratories, H-1000).

Images of eye disc tumours and whole larvae were acquired under 3.25x and 1.25x objectives respectively, using a fluorescent stereo microscope (Leica M165 FC) equipped with a digital camera (Leica DFC 3000G). Wing imaginal disc images were acquired with a confocal microscope (Leica Stellaris 5) using a 10x objective.

To quantify the GFP^+^ area in imaginal discs, a minimum of 20 animals per condition were analysed in FIJI (v2.16.0/1.54p) using a custom-made macro. Images were first converted to .tiff format and RGB channels were split. The imaginal disc area was manually defined, based on the outline of DAPI staining in the red channel. The green channel was thresholded for GFP intensity to allow for the identification of GFP^+^ cells within the user-defined imaginal disc area.

To assess ectopic tumours, a minimum of 20 larvae were scored per condition. Individual larvae were washed, and 3 images were acquired for each animal (dorsal, right lateral and left lateral views) to ensure accurate visualisation throughout the larvae. The size of each ectopic tumour was quantified by FIJI (v2.16.0/1.54p) based on GFP^+^ area relative to the total dorsal body area, and classified into the following categories: I (<0.008), II (≥0.008-<0.02), III (≥0.02-<0.035) and IV (≥0.035) (see Fig. S5G).

### Midgut immunostaining and pH3^+^ scoring

For phospho-histone-H3 (pH3) staining, adult females (*esg^ts^* driver) induced for 7 days at 29°C were anesthetised on ice. Midguts were dissected in 1X PBS and mounted on poly-L-lysine coated slides (Sigma, P1524). Tissues were fixed in 4% formaldehyde (bioWORLD, #30450002) in PBS for 30 min, washed with a wash protocol of 3 rinses with 1X PBS, 15 min wash with mild shaking, and another 3 rinses with 1X PBS, incubated in PBS-T (0.5% Triton X-100 in 1X PBS) for 20 min at RT to permeabilise the tissue, then blocked for 2 h with PBS-T + 10% normal goat serum (NGS; Abcam, AB7481) at RT with mild shaking. Samples were stained with anti-pH3 primary antibody (Ser10, 1:200; Cell Signaling Technology, #9701) overnight at 4°C in PBS-T + 10% NGS. After a wash protocol with PBS-T, slides were stained with secondary antibody (Alexa Fluor 568 goat anti-rabbit IgG; Thermo Fisher Scientific, A-11036) in PBS-T for 2 h at RT with mild shaking, then washed and mounted in Vectashield with DAPI (Vector Laboratories, H-1200). pH3^+^ cells were visually counted on a fluorescent stereomicroscope (Leica DM4 B) at 20x magnification along the entire midgut (pH3^+^ cells in the Malpighian tubules were excluded). Typical pH3^+^ images were taken for the R4 region of the midgut on a confocal microscope (Leica Stellaris 5).

### Hindgut imaging by micro-computed tomography (μCT)

To assess hindgut size and morphology *in situ* by µCT^51,52^, induced adult flies (∼20 per condition) were anesthetised on ice and transferred to a 1.5 mL tube containing 1 mL 1X PBS (Thermo, 70011044) plus 0.5% v/v Triton X-100 (Sigma, T9284) for ∼5 min. Flies were fixed in 1 mL of Bouin’s solution (Sigma, HT10132) for 24 h at RT, then washed with 1X PBS (3x 1 mL) followed by 1x 1 mL of 1X PBS each day for 3 subsequent days, stained in 1:1 of Lugol solution (Sigma, 62650):ultrapure H2O (Invitrogen, 10977035) for 72 h, and finally rinsed 1x in 1 mL ultrapure H2O. Flies were scanned in groups of 20 flies, by inserting sequentially into p10 pipette tips containing ultrapure H2O (10 flies per tip), and sealing the openings with parafilm to prevent drying out during scanning. The 2x p10 tips, each containing 10 flies, were secured tightly together with parafilm and scanned on the µCT (Bruker Skyscan 1272) with the following settings (40 kV; 110 μA; 2.95 μm pixel size; rotation step, 0.150°; averaging of 4 frames, 30 px of random movement) performed as a standard scan in a central camera position. Flatfield corrections were applied before every scan to maximise image resolution. Reconstruction was performed with Bruker NRecon software (v2.2.0.6) and GPUReconServer (v2.2.0.0), and subsequently processed in FIJI (v2.16.0/1.54p) using a custom macro^52^ to correct for scanning inversions/misalignments, adjust the contrast, and permit background subtraction to generate the final .tiff files for analysis. Whole hindgut segmentation was performed with ITK-SNAP (v4.2.0-rc.1)^74^ at 3.0x zoom. Circumferences were outlined manually every ∼2-4 slices using the polygon tool, primarily along the axial plane, followed by the interpolation tool to fully reconstruct each anterior hindgut and extract the volume. The anterior hindgut was defined as the region posterior to the junction of the Malpighian tubules and anterior to the rectal ampulla. 3D segmentations were smoothened with the ITK-SNAP Label Smoothing Tool with settings: Standard deviation of 2 µm across xyz planes.

### Dissemination assay

To assess dissemination post-induction, flies were anaesthetised on ice, and placed dorsal side-down over a ∼4 mm-wide strip of petroleum jelly (Vaseline) on a glass microscope slide. The wings and legs were tucked into the petroleum jelly to reveal the abdominal region more clearly. Dissections were performed with flies submerged in ice-cold 1X PBS. The ventral side of the abdominal cuticle was cut open along the sternites using fine scissors (Fine Science Tools, Vannas Spring Scissors straight 2.5 mm cutting edge, #15000-08) and folded over into the petroleum jelly. The exposed internal abdominal organs (i.e. reproductive, digestive and excretory) were carefully removed without disturbing the dorsal fat tissue layer. The number of GFP^+^ foci per fly abdominal cavity was counted while accounting for the z plane in a dark environment using a fluorescent stereomicroscope (Leica M165FC) at 12x magnification with a mounted 0.5x stereo C-mount objective (Leica #10450528). All dissections and scoring were carried out by the same individual with genotypes blinded.

### EdU incorporation assay

Flies were maintained at 18°C for 7 days following eclosion to allow maturation, then *UAS-Ras^G12X^* transgene expression was induced for 14 days under control of the hindgut-specific driver *byn-GAL4^ts^* by transferring flies to 29°C. During the last 7 days, flies were fed standard SYA medium supplemented with 5 mg/mL EdU. Fly hindguts were dissected on ice and fixed in 4% PFA for 30 min at RT with mild shaking (47 rpm, Stuart Mini Orbital Shaker - SSM1) and placed on poly-L-lysine-coated microscope slides, then washed with a protocol of 3x rinses with 1X PBS, 15 min wash with mild shaking, and another 3x rinses with 1X PBS. Permeabilisation was performed with 1X PBS, 1% Triton X-100 for 30 min at RT with mild shaking. Fly hindguts were then rinsed twice with 1X PBS, then incubated in Click-iT EdU Alexa Fluor 647 imaging kit cocktail (Invitrogen, C10340) for 30 min in a dark environment at RT with mild shaking. Following another wash protocol, the guts were mounted in Vectashield Antifade Mounting Medium with DAPI (Vector Laboratories, H-1200), and imaged with a 10x objective on a confocal microscope (Leica Stellaris 5). Quantification of relative EdU areas was performed with QuPath (v0.5.0-x64).

### Analysis of KRAS^G12X^ mutations in human cancers (COSMIC database)

Information on point mutations at the G12 and G13 positions in the KRAS gene (COSG97004) was obtained from the COSMIC (Catalogue of Somatic Mutations in Cancer, https://cancer.sanger.ac.uk/cosm) database (v103, released 18/11/2025, accessed 04/05/2026), which comprises 302,240 total unique samples^6^. From the 253,224 samples tested, 49,216 samples contained a mutation in KRAS, with 48,777 corresponding to a single nucleotide missense substitution (Table S1). To plot the distribution of KRAS^G12X^ mutations across tumour types, tissues containing >10% mutations were considered (Tables S3,S4)^6,7^.

### Sequence alignment and structural prediction

Protein sequence alignment between human KRAS (isoform B, UniProt: P01116-2) and *Drosophila* Ras (UniProt: P08646) was performed in Clustal Omega (v1.2.4)^75^. The structure of *Drosophila* Ras was predicted by AlphaFold 3 (accessed on 24/08/2026)^76^, using the *Drosophila* Ras protein sequence (UniProt: P08646) together with GDP and a Mg^2+^ ion as input. The predicted model showed overall confidence (pLDDT, predicted local distance difference test: >90). To compare human KRAS (PDB: 4OBE) with AlphaFold 3-generated *Drosophila* Ras, a single protein chain (residues 1-169) from each structure was super-imposed in PyMOL (v2.3.5), and the RMSD (root mean square deviation) value was obtained.

## Statistical analysis

The majority of data was plotted and analysed in GraphPad Prism (v10). Data are displayed as boxplots (n<15) or violin plots (n≥15). Lifespan assays are plotted as cumulative survival curves, with statistical analysis performed by Log-Rank test in Microsoft Excel using a custom macro. Full details of survival data (exact n numbers, p values) are provided in Table S6. Other data were analysed by one-way ANOVA or non-parametric testing, as appropriate. Statistical significance was determined as: n/s, p>0.05; *, p<0.05; **, p<0.01; ***, p<0.001.

## Data availability

This study includes no data deposited in external repositories. All data are provided within the Figures and Supplementary Information.

## Author contributions

Conceptualisation, SH and HMC; methodology, SH and HMC; investigation and formal analysis, MRV, MAT, NN, MBA, IL, AF, SH and HMC; funding acquisition, SH and HMC; supervision, SH and HMC; writing-original draft, SH and HMC, with input from all the authors.

## Acknowledgements

We are grateful to Cathie Pfleger for helpful discussions. We acknowledge Linda Partridge, Erdem Bangi, Ross Cagan, Bruce Edgar, Georg Halder, Alana O’Reilly and Michael Simon for kindly providing fly stocks, as well as the Bloomington *Drosophila* Stock Center (BDSC) (NIH P40OD018537). We thank current and former members of the Hirabayashi and Cochemé groups, particularly Cecilia Cabrera for generating the fly gut images, Emily Strachan for advice on fluorescence microscopy analysis, Laura Camplese and Mary O’Sullivan for technical support with media preparation, and Oleh Lushchak for initial training on µCT imaging. We acknowledge Dirk Dormann and Chad Whilding from the LMS Microscopy Facility for assistance with image quantification. This work was funded by the Medical Research Council UK (grants MC-M302-M3450 to HMC and MC-M302-M3510 to SH). NN is supported by a Career Development Fellowship from the Royal Society (CDF\R1\241963). For the purpose of open access, the authors have applied a Creative Commons Attribution (CC BY) licence to any Author Accepted Manuscript version arising.

## Disclosure and competing interests statement

The authors declare that they have no conflict of interest.

## Supplementary Information

### Supplementary Tables

**Table S1.**
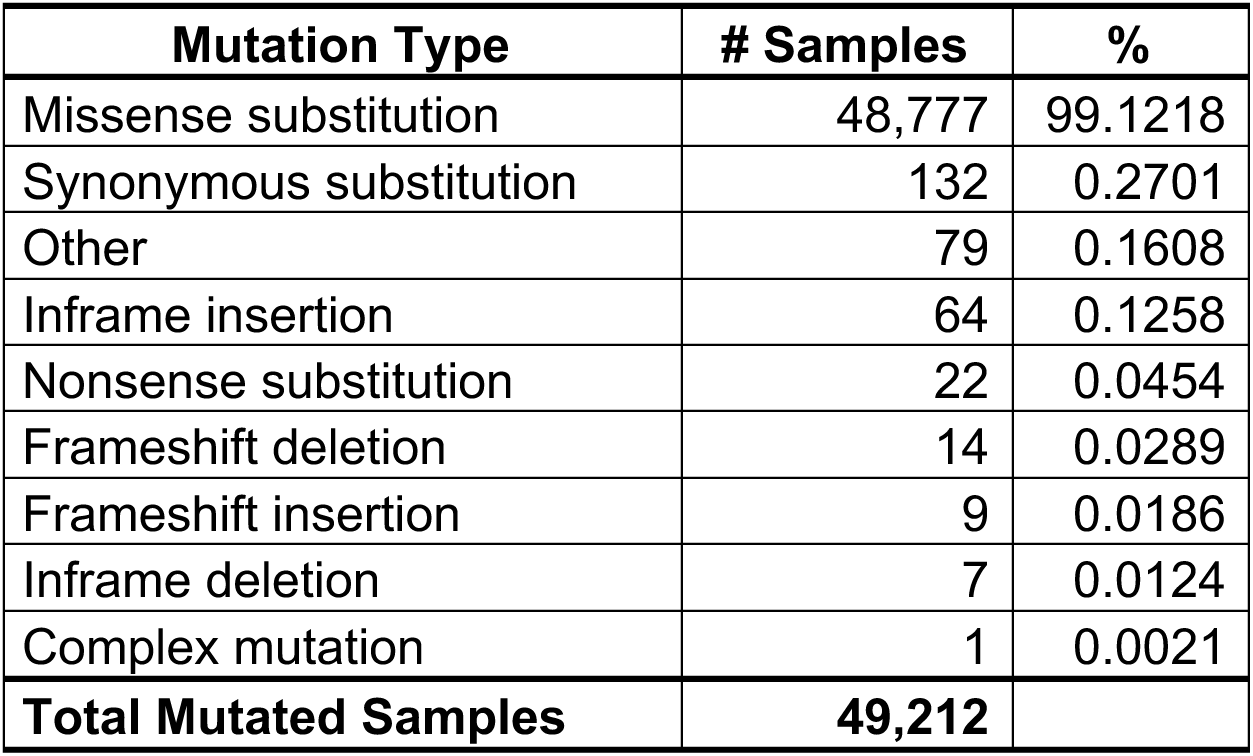
Summary of KRAS mutations in human cancers. Mutations in KRAS (COSG97004) were observed in 49,212 unique samples from a total of 253,224 tested. Based on the COSMIC database (v103, released 18/11/2025).

| Mutation Type | # Samples | % |
|---|---|---|
| Missense substitution | 48,777 | 99.1218 |
| Synonymous substitution | 132 | 0.2701 |
| Other | 79 | 0.1608 |
| Inframe insertion | 64 | 0.1258 |
| Nonsense substitution | 22 | 0.0454 |
| Frameshift deletion | 14 | 0.0289 |
| Frameshift insertion | 9 | 0.0186 |
| Inframe deletion | 7 | 0.0124 |
| Complex mutation | 1 | 0.0021 |
| **Total Mutated Samples** | **49,212** |  |

**Table S2.**
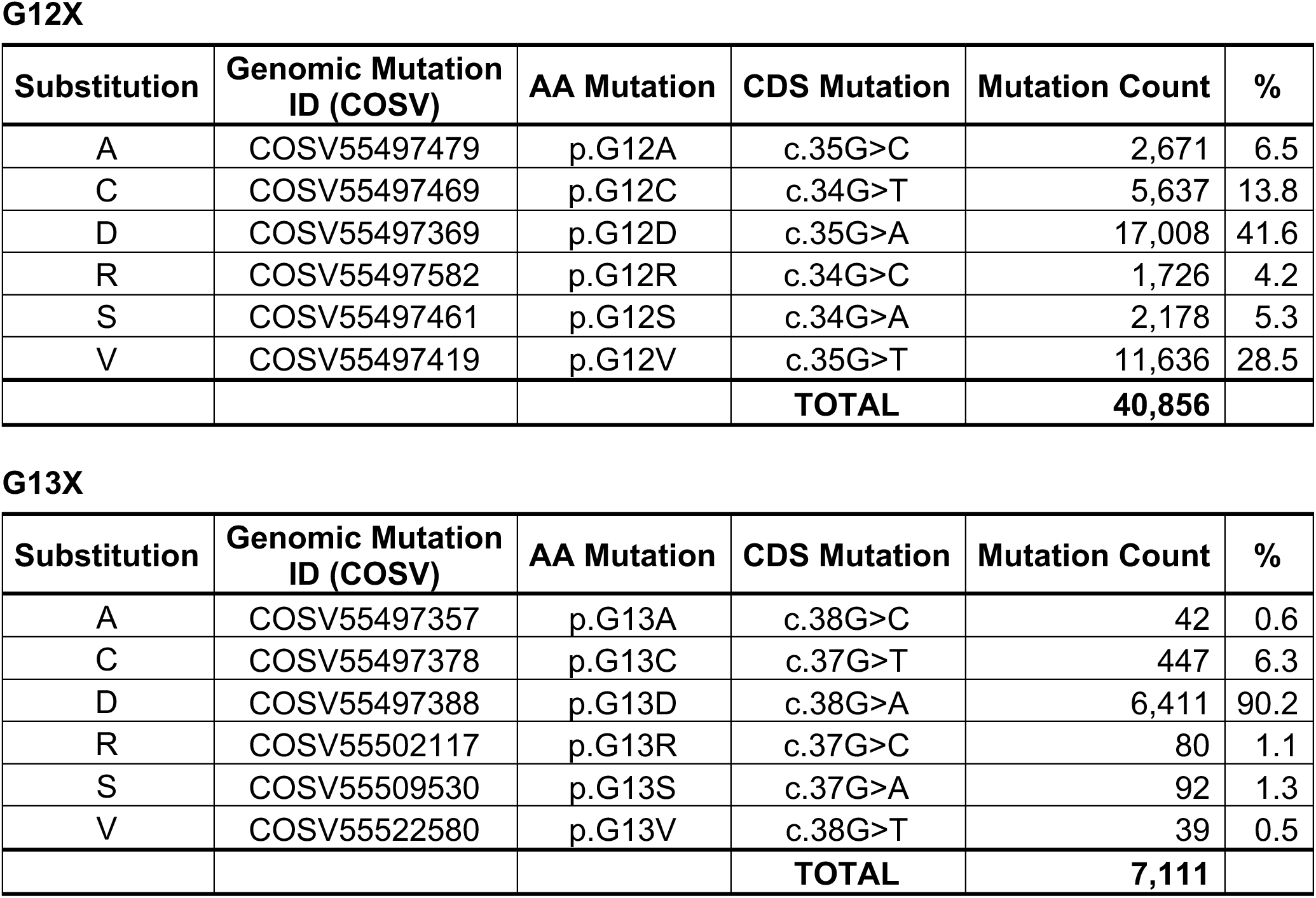
Distribution of G12X and G13X KRAS point mutations in human cancers. Missense substitutions in KRAS (COSG97004) at positions G12X and G13X across all tissues. Based on the COSMIC database (v103, released 18/11/2025).

| **G12X** |  |  |  |  |  |
|---|---|---|---|---|---|
| **Substitution** | **Genomic Mutation ID (COSV)** | **AA Mutation** | **CDS Mutation** | **Mutation Count** | **%** |
|---|---|---|---|---|---|
| A | COSV55497479 | p.G12A | c.35G>C | 2,671 | 6.5 |
| C | COSV55497469 | p.G12C | c.34G>T | 5,637 | 13.8 |
| D | COSV55497369 | p.G12D | c.35G>A | 17,008 | 41.6 |
| R | COSV55497582 | p.G12R | c.34G>C | 1,726 | 4.2 |
| S | COSV55497461 | p.G12S | c.34G>A | 2,178 | 5.3 |
| V | COSV55497419 | p.G12V | c.35G>T | 11,636 | 28.5 |
|  |  |  | **TOTAL** | **40,856** |  |
| **G13X** |  |  |  |  |  |
| **Substitution** | **Genomic Mutation ID (COSV)** | **AA Mutation** | **CDS Mutation** | **Mutation Count** | **%** |
| A | COSV55497357 | p.G13A | c.38G>C | 42 | 0.6 |
| C | COSV55497378 | p.G13C | c.37G>T | 447 | 6.3 |
| D | COSV55497388 | p.G13D | c.38G>A | 6,411 | 90.2 |
| R | COSV55502117 | p.G13R | c.37G>C | 80 | 1.1 |
| S | COSV55509530 | p.G13S | c.37G>A | 92 | 1.3 |
| V | COSV55522580 | p.G13V | c.38G>T | 39 | 0.5 |
|  |  |  | **TOTAL** | **7,111** |  |

**Table S3.** Distribution of mutations in KRAS by cancer tissue. Mutations in KRAS (COSG97004) across primary tissue types. Tissues with >10% mutated samples are listed. Based on the COSMIC database (v103, released 18/11/2025).

| Tissue | Total Mutated Samples | Total Samples Tested | % |
|---|---|---|---|
| Pancreas | 7,789 | 14,664 | 53.12 |
| Peritoneum | 167 | 372 | 44.89 |
| Large intestine | 26,695 | 83,256 | 32.06 |
| Small intestine | 311 | 1,431 | 21.73 |
| Biliary tract | 1,063 | 6,092 | 17.45 |
| Endometrium | 789 | 5,096 | 15.48 |
| Lung | 6,851 | 48,478 | 14.13 |
| Ovary | 1,135 | 8,037 | 14.12 |

**Table S4.**
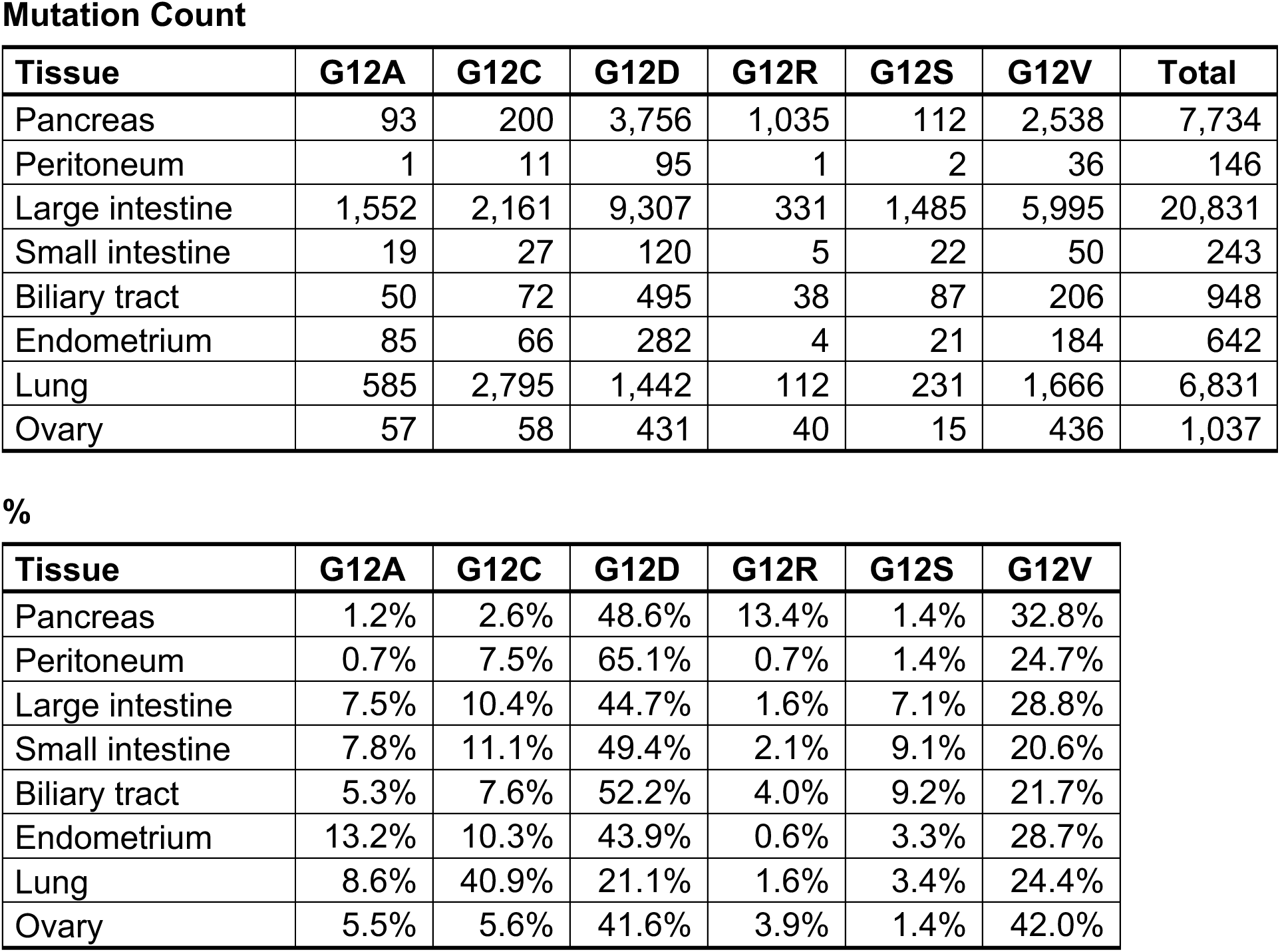
Distribution of G12X point mutations in KRAS by cancer tissue. Missense substitutions in KRAS (COSG97004) at position G12X across primary tissue types. Based on the COSMIC database (v103, released 18/11/2025).

| **Mutation Count** |  |  |  |  |  |  |  |
|---|---|---|---|---|---|---|---|
| **Tissue** | **G12A** | **G12C** | **G12D** | **G12R** | **G12S** | **G12V** | **Total** |
| Pancreas | 93 | 200 | 3,756 | 1,035 | 112 | 2,538 | 7,734 |
| Peritoneum | 1 | 11 | 95 | 1 | 2 | 36 | 146 |
| Large intestine | 1,552 | 2,161 | 9,307 | 331 | 1,485 | 5,995 | 20,831 |
| Small intestine | 19 | 27 | 120 | 5 | 22 | 50 | 243 |
| Biliary tract | 50 | 72 | 495 | 38 | 87 | 206 | 948 |
| Endometrium | 85 | 66 | 282 | 4 | 21 | 184 | 642 |
| Lung | 585 | 2,795 | 1,442 | 112 | 231 | 1,666 | 6,831 |
| Ovary | 57 | 58 | 431 | 40 | 15 | 436 | 1,037 |
| **%** |  |  |  |  |  |  |  |
| **Tissue** | **G12A** | **G12C** | **G12D** | **G12R** | **G12S** | **G12V** |  |
| Pancreas | 1.2% | 2.6% | 48.6% | 13.4% | 1.4% | 32.8% |  |
| Peritoneum | 0.7% | 7.5% | 65.1% | 0.7% | 1.4% | 24.7% |  |
| Large intestine | 7.5% | 10.4% | 44.7% | 1.6% | 7.1% | 28.8% |  |
| Small intestine | 7.8% | 11.1% | 49.4% | 2.1% | 9.1% | 20.6% |  |
| Biliary tract | 5.3% | 7.6% | 52.2% | 4.0% | 9.2% | 21.7% |  |
| Endometrium | 13.2% | 10.3% | 43.9% | 0.6% | 3.3% | 28.7% |  |
| Lung | 8.6% | 40.9% | 21.1% | 1.6% | 3.4% | 24.4% |  |
| Ovary | 5.5% | 5.6% | 41.6% | 3.9% | 1.4% | 42.0% |  |

**Table S5.**
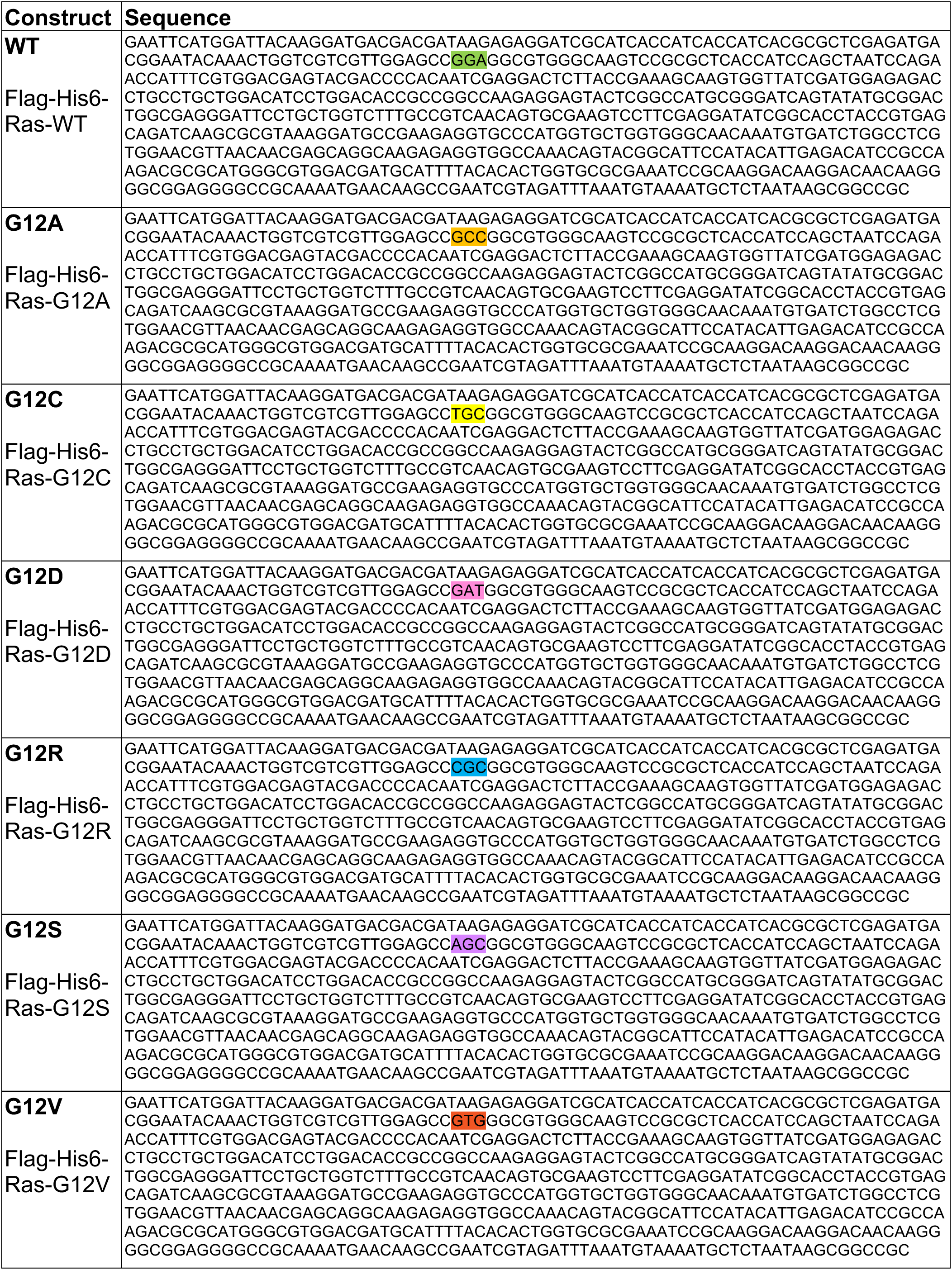
Sequences of the *UAS-Ras^G12X^* transgenes. Sequences of the N-terminally Flag- and His-tagged constructs inserted into the pUASTattB vector, used to generate the *UAS-Ras^G12X^*toolkit. The WT and mutated codons at position 12 are highlighted.

**Table S6.** Summary of survival data. Female flies were induced at d7 of adulthood by transferring vials from 18°C to 29°C. Survival post-induction was scored at 29°C.

| Figure | Genotype | Flies per vial | Vials # | Total flies (set-up) | Deaths # | Censors # | Total flies (actual) | Median survival (d) | p value (Log-Rank test) |  |  |  |  |  |  |  |  |
|---|---|---|---|---|---|---|---|---|---|---|---|---|---|---|---|---|---|
|  |  |  |  |  |  |  |  |  |  | *b* | *c* | *d* | *e* | *f* | *g* | *h* | *i* |
| **Fig. 6E &<br>Fig. S6C** | *esg*<sup>*ts*</sup>>*w*<sup>*1118*</sup> | 20 | 10 | 200 | 198 | 0 | 198 | 45.43 | *a* | 5.0E-05 | 5.3E-96 | 1.0E-95 | 3.3E-97 | 4.4E-72 | 2.7E-98 | 4.5E-103 | 1.2E-98 |
|  | *esg*<sup>*ts*</sup>>*Ras*<sup>*WT*</sup> | 20 | 10 | 200 | 200 | 1 | 201 | 46.72 | *b* |  | 1.6E-96 | 2.3E-96 | 1.5E-98 | 6.8E-75 | 7.4E-99 | 8.7E-104 | 3.1E-99 |
|  | *esg*<sup>*ts*</sup>>*Ras*<sup>*G12A*</sup> | 20 | 10 | 200 | 196 | 0 | 196 | 29.24 | *c* |  |  | 0.5201 | 0.0002 | 1.7E-47 | 0.0003 | 1.0E-26 | 2.0E-06 |
|  | *esg*<sup>*ts*</sup>>*Ras*<sup>*G12C*</sup> | 20 | 10 | 200 | 199 | 0 | 199 | 28.66 | *d* |  |  |  | 0.0033 | 7.9E-43 | 0.0238 | 2.2E-14 | 0.0005 |
|  | *esg*<sup>*ts*</sup>>*Ras*<sup>*G12D*</sup> | 20 | 10 | 200 | 200 | 1 | 201 | 23.80 | *e* |  |  |  |  | 2.3E-50 | 0.2618 | 0.0001 | 0.9491 |
|  | *esg*<sup>*ts*</sup>>*Ras*<sup>*G12R*</sup> | 20 | 10 | 200 | 198 | 0 | 198 | 36.45 | *f* |  |  |  |  |  | 2.0E-59 | 1.2E-79 | 7.6E-64 |
|  | *esg*<sup>*ts*</sup>>*Ras*<sup>*G12S*</sup> | 20 | 10 | 200 | 200 | 0 | 200 | 27.53 | *g* |  |  |  |  |  |  | 1.6E-12 | 0.1747 |
|  | *esg*<sup>*ts*</sup>>*Ras*<sup>*G12V*</sup> | 20 | 10 | 200 | 198 | 1 | 199 | 20.29 | *h* |  |  |  |  |  |  |  | 3.5E-07 |
|  | *esg*<sup>*ts*</sup>>*Ras*<sup>*G12V(P)*</sup> | 20 | 10 | 200 | 198 | 1 | 199 | 26.46 | *i* |  |  |  |  |  |  |  |  |
|  |  |  |  |  |  |  |  |  |  | *b* | *c* | *d* | *e* | *f* | *g* | *h* | *i* |
| **Fig. 7I &<br>Fig. S7G** | *byn*<sup>*ts*</sup>>*w*<sup>*1118*</sup> | 20 | 7 | 140 | 135 | 1 | 136 | 45.95 | *a* | 0.2622 | 5.1E-61 | 3.4E-58 | 1.1E-67 | 6.7E-33 | 1.2E-71 | 1.0E-56 | 3.3E-67 |
|  | *byn*<sup>*ts*</sup>>*Ras*<sup>*WT*</sup> | 20 | 10 | 200 | 195 | 2 | 197 | 45.56 | *b* |  | 9.1E-72 | 8.1E-69 | 2.0E-83 | 5.4E-33 | 8.5E-91 | 1.7E-67 | 5.9E-82 |
|  | *byn*<sup>*ts*</sup>>*Ras*<sup>*G12A*</sup> | 20 | 10 | 200 | 202 | 0 | 202 | 35.91 | *c* |  |  | 0.0982 | 1.4E-09 | 7.1E-19 | 3.2E-21 | 0.0343 | 0.0002 |
|  | *byn*<sup>*ts*</sup>>*Ras*<sup>*G12C*</sup> | 20 | 9 | 180 | 178 | 1 | 179 | 34.24 | *d* |  |  |  | 0.0002 | 2.1E-19 | 2.7E-11 | 0.7499 | 0.1372 |
|  | *byn*<sup>*ts*</sup>>*Ras*<sup>*G12D*</sup> | 20 | 9 | 180 | 179 | 0 | 179 | 32.79 | *e* |  |  |  |  | 3.8E-36 | 0.0011 | 0.0007 | 0.0050 |
|  | *byn*<sup>*ts*</sup>>*Ras*<sup>*G12R*</sup> | 20 | 8 | 160 | 157 | 0 | 157 | 40.21 | *f* |  |  |  |  |  | 4.1E-50 | 1.8E-20 | 3.5E-31 |
|  | *byn*<sup>*ts*</sup>>*Ras*<sup>*G12S*</sup> | 20 | 10 | 200 | 200 | 2 | 202 | 31.68 | *g* |  |  |  |  |  |  | 5.7E-10 | 1.5E-10 |
|  | *byn*<sup>*ts*</sup>>*Ras*<sup>*G12V*</sup> | 20 | 8 | 160 | 148 | 3 | 151 | 34.43 | *h* |  |  |  |  |  |  |  | 0.2971 |
|  | *byn*<sup>*ts*</sup>>*Ras*<sup>*G12V(P)*</sup> | 20 | 10 | 200 | 197 | 1 | 198 | 34.38 | *i* |  |  |  |  |  |  |  |  |

### Supplementary Figure Legends

**Figure S1.**
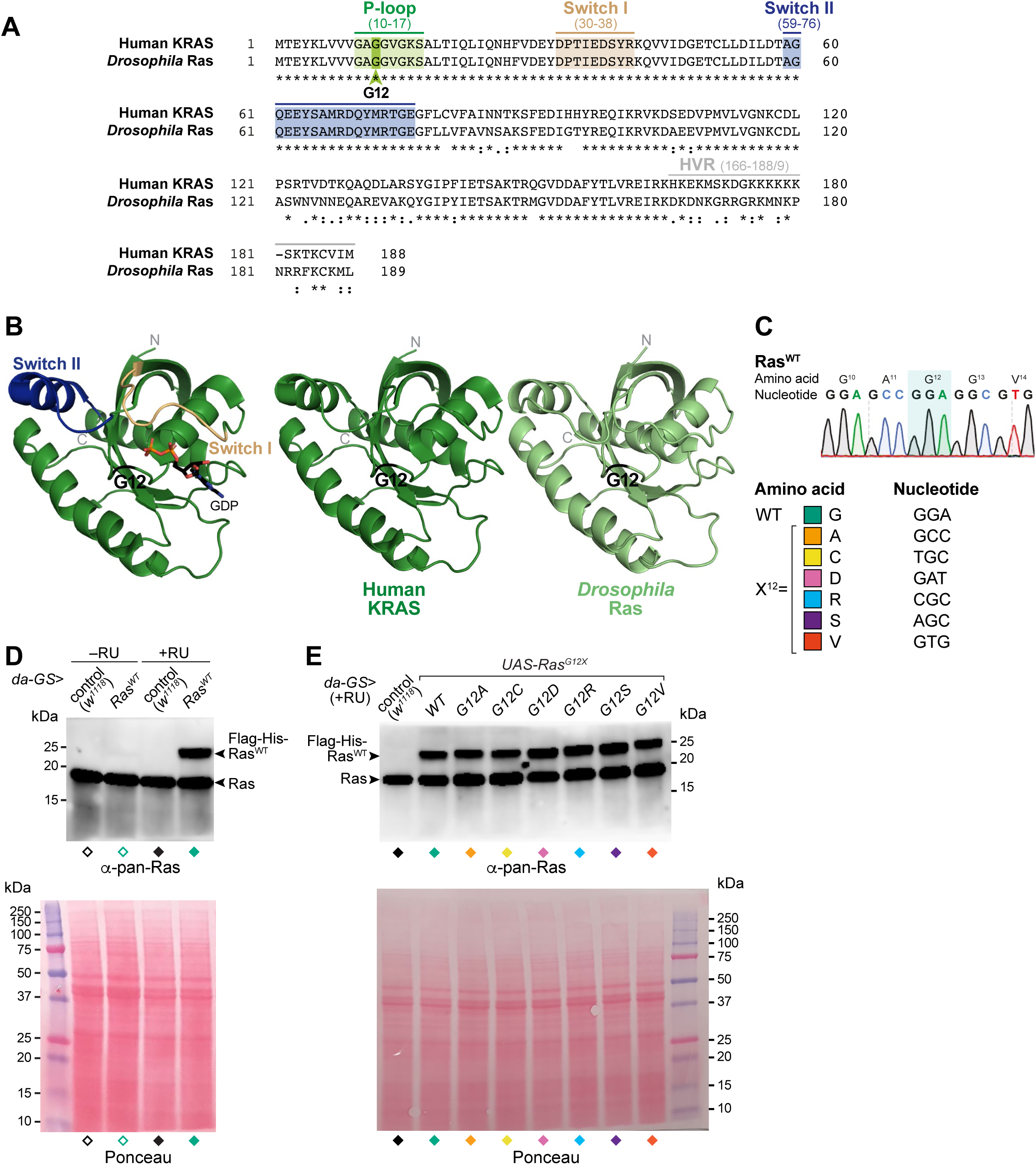
Generation and validation of the Ras^G12X^ *Drosophila* toolkit. (**A**) Protein sequence alignment between human KRAS (isoform B; UniProt: P01116-2) and *Drosophila* Ras (UniProt: P08646). The full protein is ∼79.8% identical, with ∼93.3% identity across residues 1-120. Notably, the first N-terminal 79 amino acids are 100% identical between flies and humans, which includes several important functional domains: the ‘P-loop’ motif (10-17, responsible for phosphate binding), the ‘Switch I’ region (30-38, forming the primary binding interface with downstream effector proteins such as RAF and PI3K, as well regulators such as GAPs and GEFs), and the ‘Switch II’ region (59-76, containing a conserved catalytic glutamine, Q61, required for GTP hydrolysis). The last C-terminal ∼20 residues correspond to the hypervariable region (HVR), known to be poorly conserved between isoforms, which dictates membrane localisation and cellular trafficking. (**B**) Left: Protein structure of human KRAS (isoform B; PDB: 4OBE, residues 1-169), highlighting the ‘Switch I’ and ‘Switch II’ regions in beige and dark blue, respectively. GDP is shown in stick conformation and the magnesium ion (Mg^2+^) as a brown sphere. Right: Structure of human KRAS and AlphaFold 3 model of *Drosophila* Ras (residues 1-169). G12 is indicated in black. See Fig. 1D for the overlay. (**C**) Summary of the Ras^G12X^ point mutations and corresponding nucleotide changes introduced for the transgenic toolkit (see Table S5 for the full sequences). (**D**,**E**) Full images corresponding to the western blot data in Fig. 1G,H.

**Figure S2.**
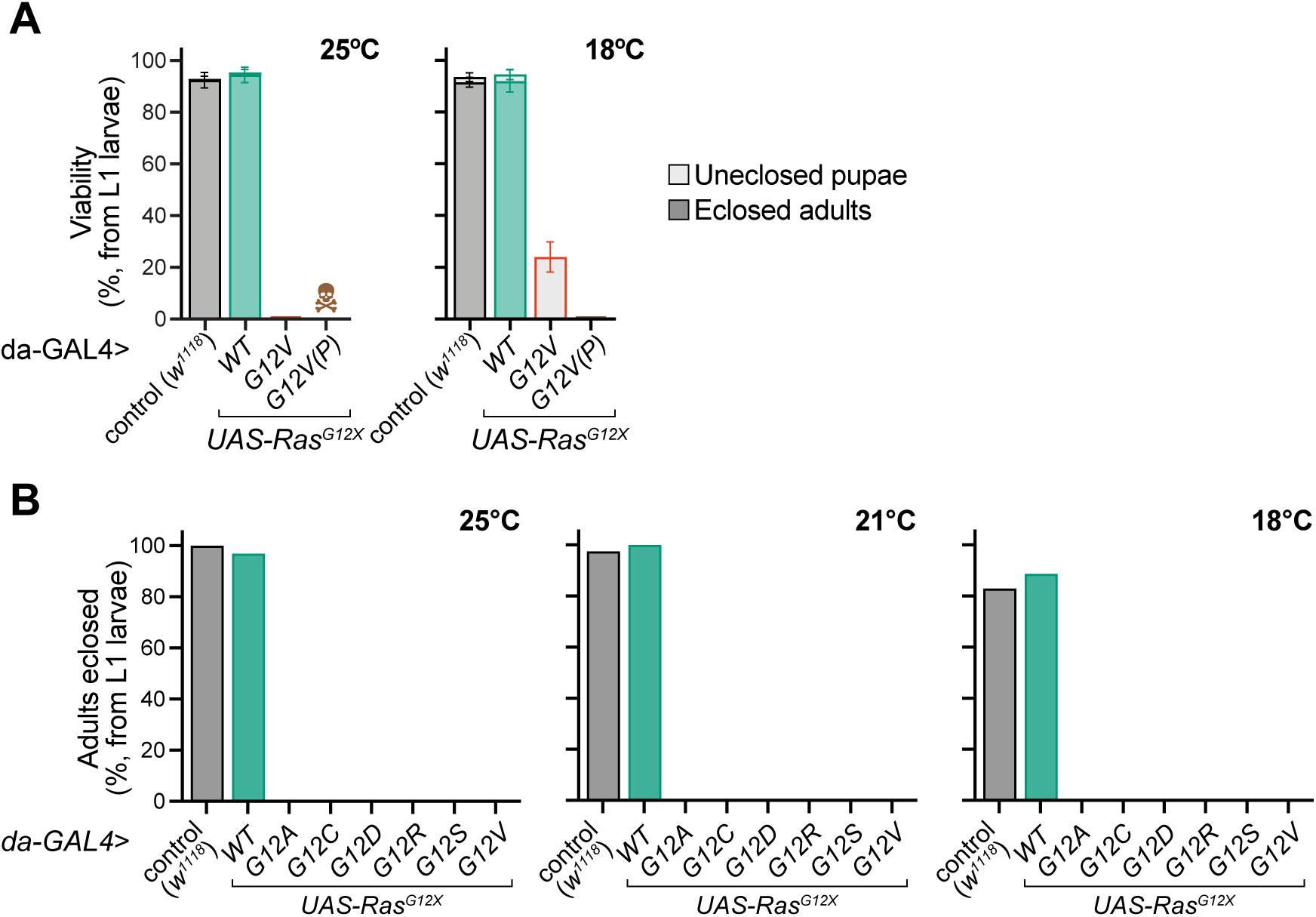
Whole body expression of Ras^G12X^ variants differentially affects development and pupariation. (**A**) Scoring of viability to pupae and eclosed adults (shaded and solid colour-coding, respectively) for the *da-GAL4* driver at 25°C and 18°C, comparing the toolkit attP40 *Ras^G12V^* variant with the P-element *Ras^G12V^*(P). At 25°C, all *da-GAL4>UAS-Ras^G12V^* larvae died at the L1 stage, whereas the *da-GAL4>UAS-Ras^G12V^*(P) combination was embryonic lethal. Data are means ±SD of n=4 vials, each with n=50 picked L1 larvae (200 total). (**B**) Quantification of adult eclosion (% of L1 larvae picked) for *da-GAL4>UAS-Ras^G12X^* flies at 25°C, 21°C and 18°C. Experiments were set up with a total of n=50 (25°C) or n=200 (21°C and 18°C) L1 larvae (related to the pupariation data from Fig. 2B).

**Figure S3.**
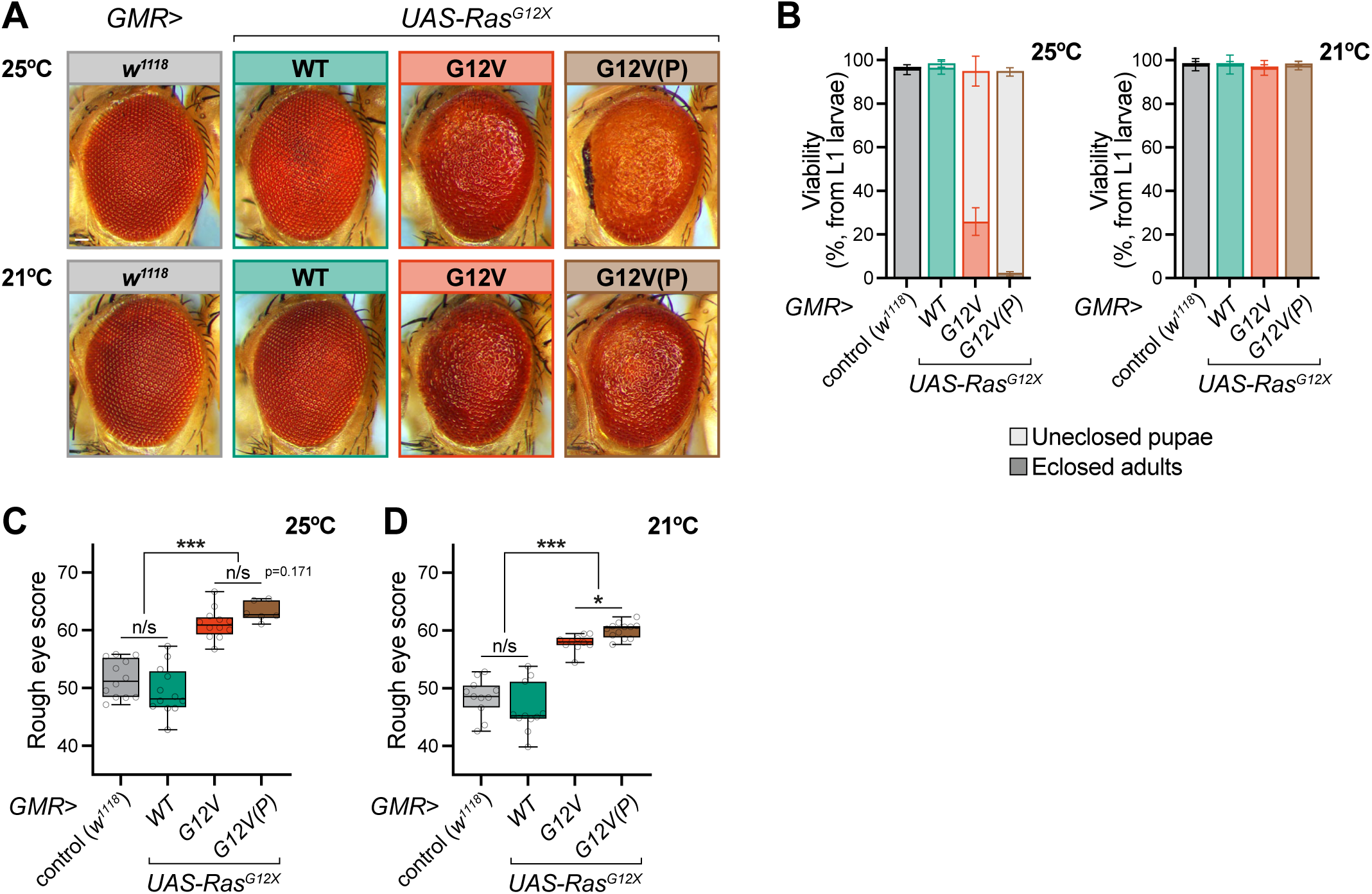
Expression of Ras^G12X^ variants in the developing eye induces differential morphological defects. (**A**) Representative z-stack images of adult female eyes, eclosed following incubation at 25°C or 21°C, comparing the toolkit attP40 *Ras^G12V^* variant with the P-element *Ras^G12V^*(P) (scale bar: 50 μm). (**B**) Scoring of viability to pupae and eclosed adults (shaded and solid colour-coding, respectively) for the *GMR-GAL4* driver, comparing the toolkit attP40 *Ras^G12V^*variant with the P-element *Ras^G12V^*(P). Data are means ±SD of n=4 vials per condition, each with n=50 picked L1 larvae (200 total). (**C**,**D**) Automated scoring of eye roughness at 25°C (**C**) and 21°C (**D**) using Flynotyper, comparing the toolkit attP40 *Ras^G12V^* variant with the P-element *Ras^G12V^*(P). Data are boxplots (line at median, interquartile range, min-to-max whiskers) of n=6-12 females per condition, analysed by one-way ANOVA. n/s, p>0.05; *, p<0.05; ***, p<0.001.

**Figure S4.**
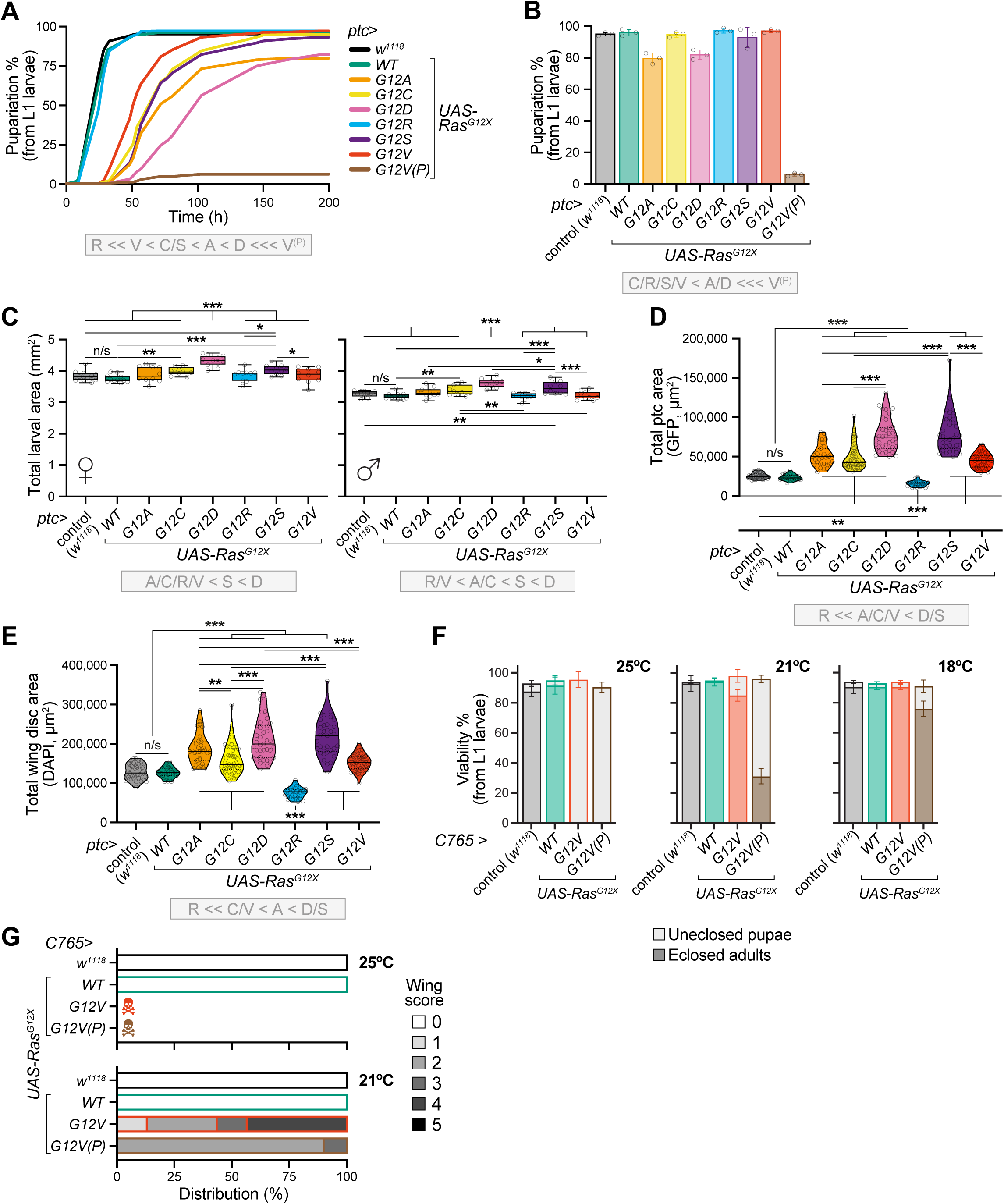
Tissue-specific expression of Ras^G12X^ variants differentially affects wing development. (**A**,**B**) Developmental phenotyping at 25°C with the *ptc-GAL4* driver, comparing the controls (*ptc*>*w*^1118^ and *ptc*>*Ras^WT^*) with the attP40 *Ras^G12X^* toolkit, and the original P-element *Ras^G12V^*(P) line. Developmental time course of pupariation (**A**) and final % pupariation for each genotype (**B**). Data are means ±SD of n=3 vials, each with n=100 picked L1 larvae (300 total). Note that the *ptc*>*Ras^G12V^*(P) combination showed strong embryonic lethality, with only a low proportion of viable L3, and was therefore not considered for further experiments. (**C**) Body size from the dorsal view area of female (left) and male (right) L3 larvae. Data are boxplots (line at median, interquartile range, min-to-max whiskers) for n=10-11 larvae per genotype, analysed by one-way ANOVA. (**D**,**E**) Absolute area of the ptc domain based on quantification of GFP (**D**) and total wing disc size measured by DAPI staining (**E**). Data are violin plots (solid line median, dotted line quartiles) of n=25-47 larvae per condition, analysed by one-way ANOVA. (**F**) Pupariation rate for the *C765-GAL4* driver at 25°C, 21°C and 18°C, comparing the toolkit attP40 *Ras^G12V^* variant with the P-element *Ras^G12V^*(P) line. Data are means ±SD of n=4 vials, each with n=50 picked L1 larvae (200 total). (**G**) Quantification of wing defects comparing the toolkit attP40 *Ras^G12V^* variant with the P-element *Ras^G12V^*(P) line. Data are bar graphs for n=6-24 flies per condition. n/s, p>0.05; *, p<0.05; **, p<0.01; ***, p<0.001.

**Figure S5.**
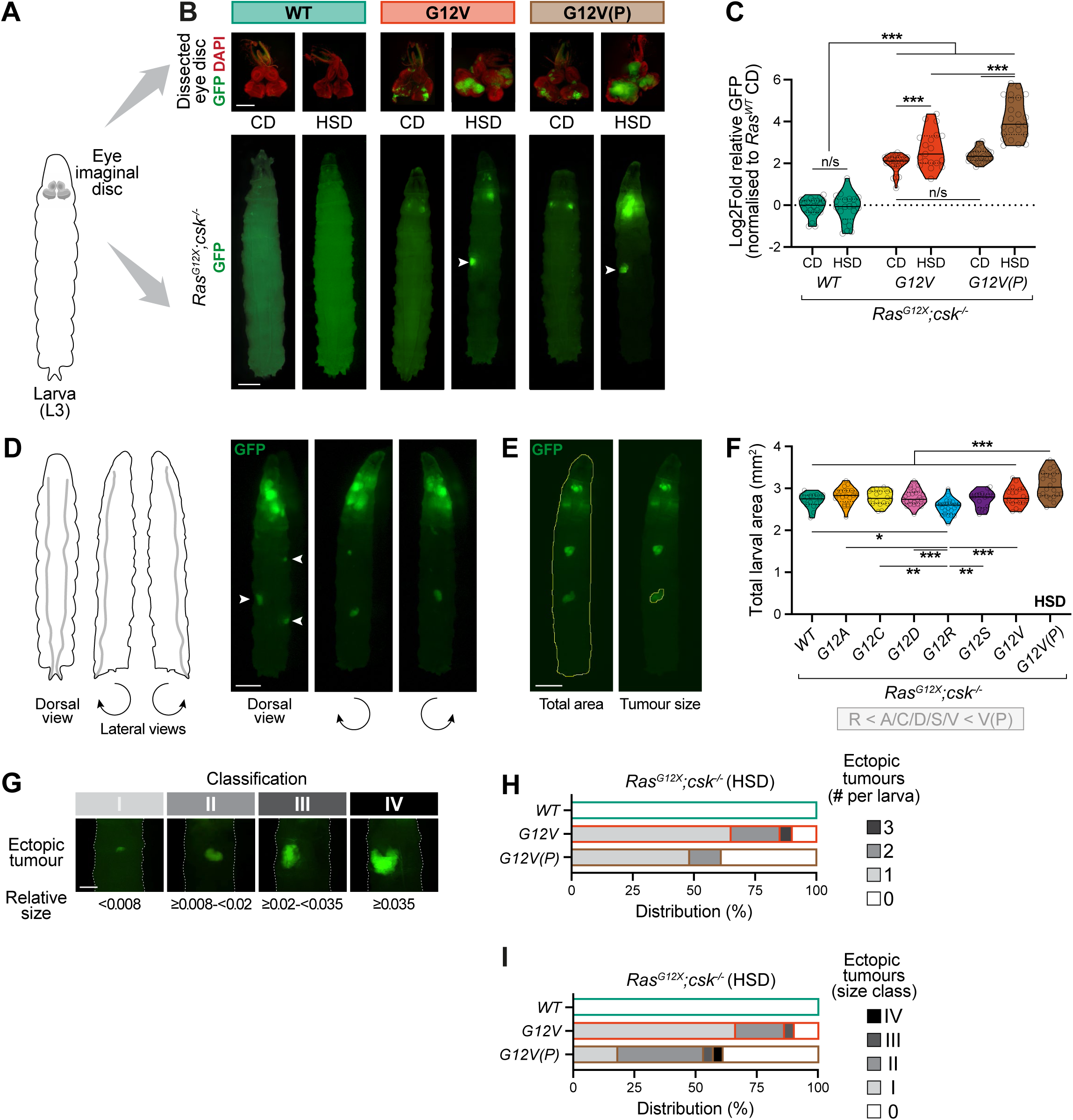
Ras^G12X^/Src co-activated eye disc clones exhibit variable responses to a high-sugar diet (HSD). (A) Diagram of an L3 larva, showing the position of the eye imaginal discs. Ras/Src co-activation in this tissue is an established model of epithelial tumour formation. (B) Representative images of developmental stage-matched L3 larvae fed a control diet (CD; day 7 after egg laying) or high-sugar diet (HSD; day 11 after egg laying), comparing the toolkit attP40 *Ras^G12V^* variant with the P-element *Ras^G12V^*(P). Above: Fluorescence microscopy of DAPI-stained dissected eye imaginal discs (red), with transformed tissue labelled by GFP (green) (scale bar: 250 μm). Below: Matched images of whole larvae (scale bar: 500 μm), with ectopic tumours observed in a subset of larvae fed a HSD (arrowheads). (C) Eye disc tumour size, based on quantification of GFP area relative to the Ras^WT^/Src control. Data are violin plots (solid line median, dotted line quartiles) of n=19-26 larvae per condition, analysed by two-way ANOVA. (**D**) Fluorescence images of an L3 larva showing the dorsal view, with ectopic tumours indicated by arrowheads, and corresponding lateral views upon rotation (scale bar: 500 μm). (**E**) Example outputs of area quantification by FIJI, showing outlining of total larval size (left), and detection of individual ectopic tumours (right) (scale bar: 500 μm). (**F**) Total larval size for the different genotypes on a HSD, from the dorsal view area (mm^2^). Data are violin plots (solid line median, dotted line quartiles) of n=15-25 larvae per condition, analysed by one-way ANOVA. (**G**) Representative images illustrating the scoring system used to classify ectopic tumour size (scale bar: 250 μm). The categories (calculated as GFP area, relative to total larval area) were defined as follows: I, <0.008; II, ≥0.008-<0.02; III, ≥0.02-<0.035; IV, ≥0.035 (related to Figs. 5F and S5H,I). (**H**,**I**) Quantification of ectopic tumour number (**H**) and size (**I**), comparing the toolkit attP40 *Ras^G12V^* variant with the P-element *Ras^G12V^*(P) line. Bar charts for n=20-25 larvae per condition. n/s, p>0.05; *, p<0.05; **, p<0.01; ***, p<0.001.

**Figure S6.**
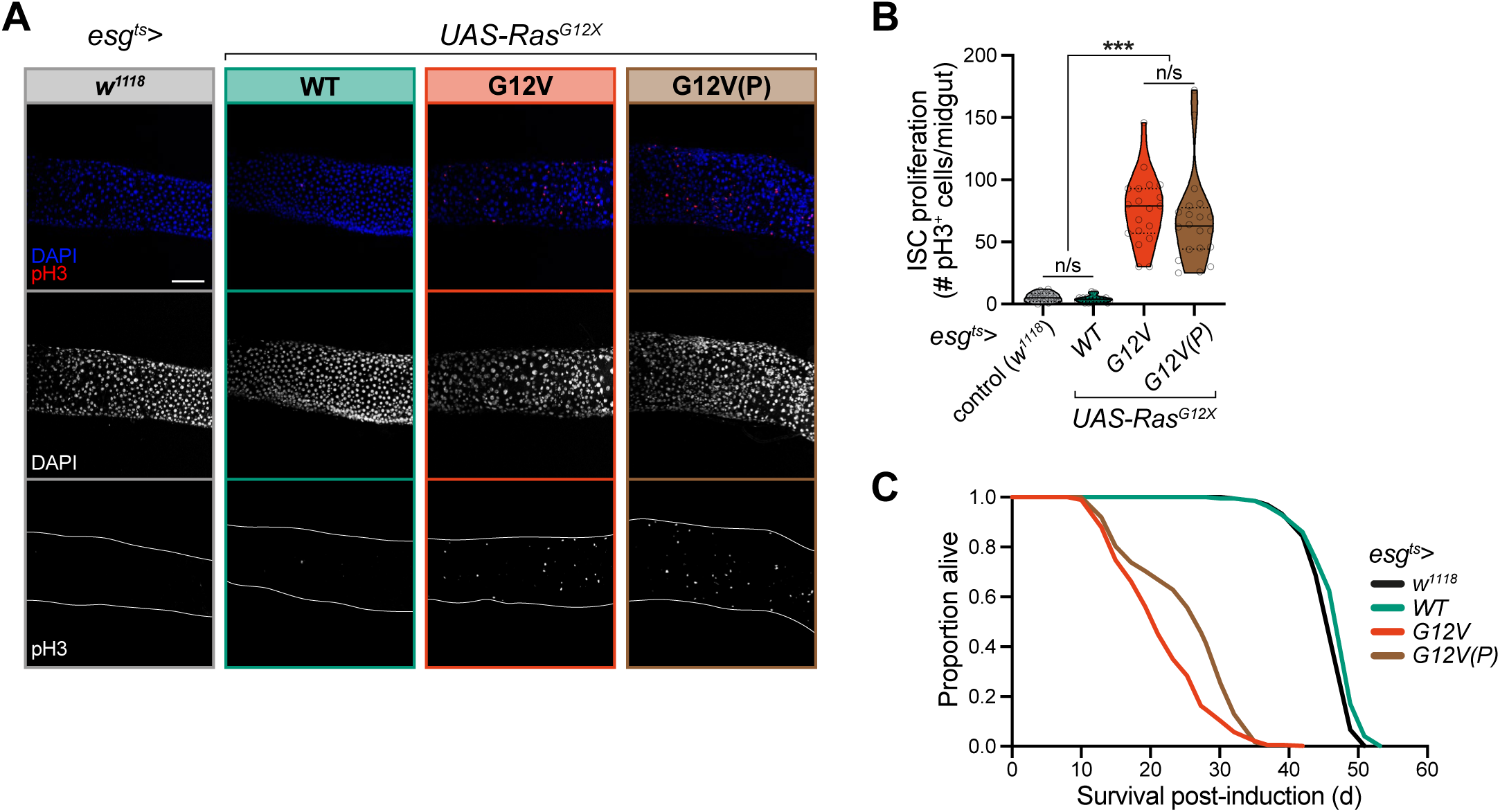
Impact of Ras^G12X^ variants on proliferation of midgut ISCs. (A) Representative confocal images of pH3^+^ cells (red), indicative of proliferation (mitosis), and DAPI staining of all nuclei (blue), comparing the toolkit attP40 *Ras^G12V^* variant with the P-element *Ras^G12V^*(P). The R4 region of the posterior midgut is shown (scale bar: 100 µm). (B) Scoring of pH3^+^ cells across the entire midgut. Data are violin plots (solid line median, dotted line quartiles) of n=19-21 females per genotype, analysed by one-way ANOVA. (C) Survival curves post-tumour induction. Lifespans were performed with n=∼200 females per condition (see Table S6 for full details of n numbers and Log-Rank p values). The pH3^+^ imaging (**A**) and scoring (**B**) were performed on controls (*esg^ts^*>*w*^1118^ and *esg^ts^*>*UAS-Ras^WT^*) and the G12V variants (*byn^ts^*>*UAS-Ras^G12V^* and *byn^ts^*>*Ras^G12V^*(P)). Females were induced at d7 of adulthood for 7 days at 29°C. n/s, p>0.05; ***, p<0.001.

**Figure S7.**
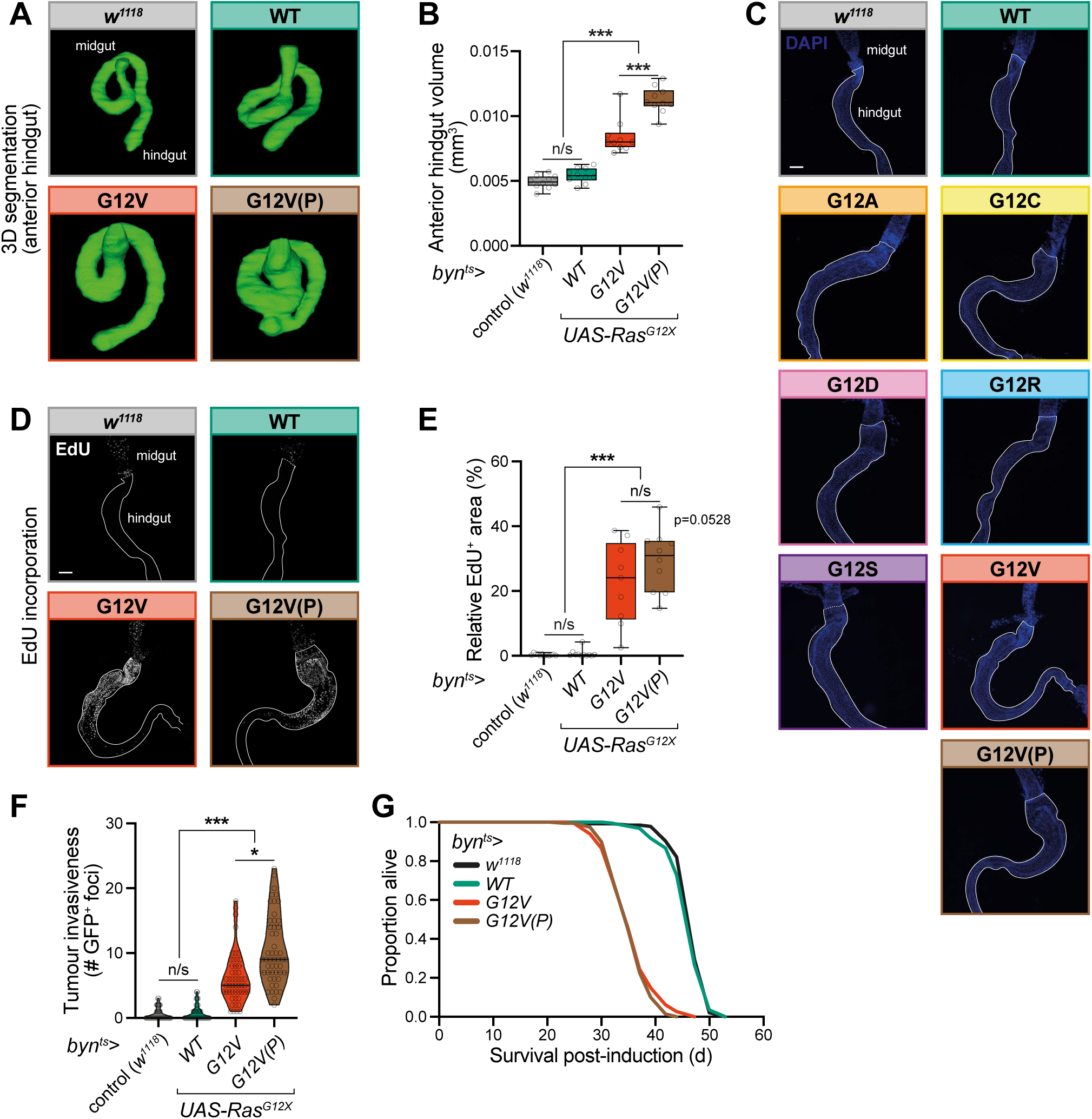
Impact of Ras^G12X^ variants in an adult colon cancer model. **A,B** µCT imaging and analysis of adult hindguts, comparing the toolkit attP40 *Ras^G12V^* variant with the P-element *Ras^G12V^*(P) line (see Figs. 7c-e). Representative 3D reconstruction (**A**) and volume quantification (**B**) of the anterior hindgut (defined as the pylorus valve and ileum, excluding the rectal ampulla). Data are boxplots (line at median, interquartile range, min-to-max whiskers) for n=10 females per genotype, analysed by one-way ANOVA. (**C**-**E**) Assessment of cell proliferation based on EdU incorporation, comparing the toolkit attP40 *Ras^G12V^* variant with the P-element *Ras^G12V^*(P) line (see Fig. 7F,G). (**C**) DAPI staining of dissected guts corresponding to Fig. 7F (scale bar: 100 µm). (**D**) Typical confocal images of hindguts, outlined by solid lines, and the midgut/hindgut junction indicated by a dotted line (scale bar: 100 µm). (**E**) Quantification of the EdU^+^ area, relative to the total anterior hindgut area. Data are boxplots (line at median, interquartile range, min-to-max whiskers) for n=9-12 flies per genotype, analysed by one-way ANOVA. (**F**) Dissemination assay data, comparing the toolkit attP40 *Ras^G12V^* variant with the P-element *Ras^G12V^*(P) line (see Fig. 7H). Data are violin plots (solid line median, dotted line quartiles) of n=50 females per genotype, analysed by a Kruskal-Wallis test. (**G**) Survival curves post-tumour induction from d7 of adulthood, comparing the toolkit attP40 *Ras^G12V^* variant with the P-element *Ras^G12V^*(P) line. Lifespans were performed with n=∼140-200 females per condition (see Table S6 for full details of n numbers and Log-Rank p values). The µCT imaging (**A,B**), EdU staining (**C-E**), and dissemination data (**F**) were performed on controls (*byn^ts^*>*w*^1118^ and *byn^ts^*>*UAS-Ras^WT^*) and the G12V variants (*byn^ts^*>*UAS-Ras^G12V^* and *byn^ts^*>*Ras^G12V^*(P)). Females were induced at d7 of adulthood for 14 days at 29°C. n/s, p>0.05; *, p<0.05; ***, p<0.001.

## Notes

### Competing Interest Statement

The authors have declared no competing interest.

